# CellTFusion: a transcriptional regulatory network framework for the identification of functional multicellular states from bulk RNA-seq data

**DOI:** 10.64898/2026.06.30.735682

**Authors:** Marcelo Hurtado, Vera Pancaldi

**Affiliations:** Université de Toulouse, INSERM, CNRS, CRCT, Toulouse, France; Equipe labellisée Ligue Contre le Cancer

## Abstract

Bulk RNA-seq remains the most accessible transcriptomic platform for tumor microenvironment (TME) characterization, yet existing computational approaches treat cell type abundance, pathway activity, and transcription factor (TF) activity as independent sources of information, missing the coordinated regulatory programs that define functional multicellular states. Here we introduce CellTFusion, a framework that integrates cell type deconvolution with transcriptional regulatory network analysis from bulk RNA-seq data to identify functional multicellular groups. CellTFusion produces a mixture representation of the TME in which each patient is described as a weighted combination of states, each capturing a distinct coordinated hallmark program. Applied to melanoma and bladder cancer cohorts, CellTFusion identified recurrent TME programs with opposing associations with immunotherapy responses that only emerged through joint multivariate modeling, and demonstrated superior cross-cohort transferability compared to established TME characterization tools.

## Introduction

Cancer remains one of the leading causes of mortality worldwide, with treatment outcomes varying across patients even within the same cancer type and treatment regimen. A central determinant of this variability is the tumor microenvironment (TME), defined as the ecosystem surrounding tumor cells; comprising immune cells, stromal cells, endothelial cells and cancer-associated fibroblasts (CAFs). This intricate ecosystem is shaped by dynamic interactions between cancer cells and immune cells, which collectively contribute to patients’ clinical outcome, metastasis and tumor progression. The advent of immune checkpoint blockade (ICB) immunotherapy, which reactivates cytotoxic immune responses against tumor cells, has underscored the critical role of the TME in determining therapeutic outcome. While ICB has transformed the treatment of multiple cancer types, durable responses are observed only in a minority of patients, and the mechanisms underlying resistance remain incompletely understood. Predicting which patients will benefit from immunotherapy therefore requires a precise and comprehensive characterization of the TME and the functional states of its components. Advances in single-cell technologies have provided detailed insights into the cellular composition of the TME, yet their high cost and complexity limit their widespread clinical application. In contrast, bulk RNA-seq remains the most accessible and scalable transcriptomic platform available in both research and clinical settings, but its aggregated nature collapses the contributions of all cell types into a single mixed signal, complicating the extraction of actionable insights and biological interpretation. Bridging this gap, and being able to extract high-resolution functional TME information from widely available bulk data is therefore a central challenge for computational oncology.

Several computational frameworks have been developed to extract TME information from bulk RNA-seq data. Deconvolution methods, including first-generation signature based approaches and second-generation machine learning tools, estimate the relative abundance of cell types from bulk expression profiles. Tools such as immunedeconv (Sturm et al., 2019), omnideconv (Dietrich et al., 2026) and multideconv (Hurtado et al., 2025) provide a unified interface to multiple deconvolution algorithms, making estimations of cell abundance easily applicable in different cancer settings for the characterization of immune infiltration across cancer cohorts.

Beyond cell type quantification, novel frameworks or methods such as EaSIeR (Lapuente-Santana et al., 2021) and iHet (Lapuente-Santana et al., 2024) adopt a systems approach, integrating cell type fractions with pathway activity, immunescores and transcription factor (TF) scores derived from prior-knowledge gene regulatory networks to build TME descriptors predictive of ICB response or associated to patient survival (Hurtado et al., 2024). Notably, iHet demonstrated that capturing both immune activation and immune exclusion components, including fibrosis-associated TGFβ signaling and CAF abundance from H&E images, improves prediction beyond immune infiltration alone. Multicellular community approaches such as EcoTyper (Luca et al., 2021) identify co-occurring cell type states from bulk data deconvolution, while markers like PD-L1 expression provide simpler single-feature proxies for immune response. Cell-cell communication inference methods, including those based on ligand-receptor co-expression databases from OmniPath (Türei et al., 2016), add an intercellular signaling dimension that is increasingly recognized as a mechanistically important component of TME characterization for immunotherapy response (van Santvoort et al., 2025).

A critical but frequently ignored aspect of this characterization is the distinction between active immune infiltration and suppressive immune states. Many existing approaches quantify immune infiltration primarily through the abundance of cytotoxic T cells or immunescores, yet tumors with high T cell infiltration can still fail to respond to immunotherapy when immune activity is counterbalanced by suppressive programs such as TGFβ, CAF-mediated stromal barriers, and immunosuppressive myeloid cells. Methods such as iHet have begun to address this aspect by explicitly modeling immune exclusion components alongside immune activation, demonstrating that disentangling these two programs from bulk RNA-seq data significantly improves the prediction of ICB response (Lapuente-Santana et al., 2024). This distinction matters because suppressive infiltration is not the absence of immune cells but an active and targetable TME state that requires functional rather than purely compositional characterization.

Despite this progress, existing bulk methods either focus on individual feature views of the TME, such as cell type abundance alone or pathway activity alone, or combine multiple views through unsupervised dimensionality reduction methods, such as multi-omics factor analysis, without explicitly grounding them by searching for a biological concordance between cellular composition and regulatory activity. This leaves a gap between compositional and functional TME characterization that current methods do not integrate.

To address this, we present CellTFusion, a computational framework that integrates cell type deconvolution with transcriptional regulatory network (TRN) analysis from bulk RNA-seq data to identify functional cell type groups, defined by shared transcription factor activity programs. This approach captures multiple cell phenotypes and states within patient samples, enhancing the characterization of the TME.

Through application to melanoma, bladder cancer, and non-small cell lung cancer cohorts, we demonstrate that CellTFusion identifies latent TME factors predictive of immunotherapy response and patient survival, transferable across independent datasets via a projection framework, and benchmarked against state-of-the-art TME characterization tools. Together, these results establish CellTFusion as a scalable and interpretable framework for extracting functional TME profiles from widely available bulk RNA-seq data, with direct applicability to clinical biomarker discovery and patient stratification. Finally, we provide an R package to the community to facilitate TME characterization of bulk RNA-seq samples, accompanied by an interactive RShiny application enabling point-and-click code-free exploration of CellTFusion outputs for researchers without computational expertise.

## Results

### CellTFusion, a transcriptional regulatory network approach for the integration of cell type deconvolution features from bulk RNAseq data

We developed CellTFusion, a computational framework designed to infer tumor microenvironment (TME) profiles from bulk RNA-seq data by integrating cell type deconvolution with transcriptional regulatory network (TRN) analysis. CellTFusion leverages both first- and second-generation deconvolution methods together with prior-knowledge gene regulatory networks to characterize coordinated multicellular functional states from bulk transcriptomic data. An overview of the full workflow is shown in Figure 1.

**Figure 1.**
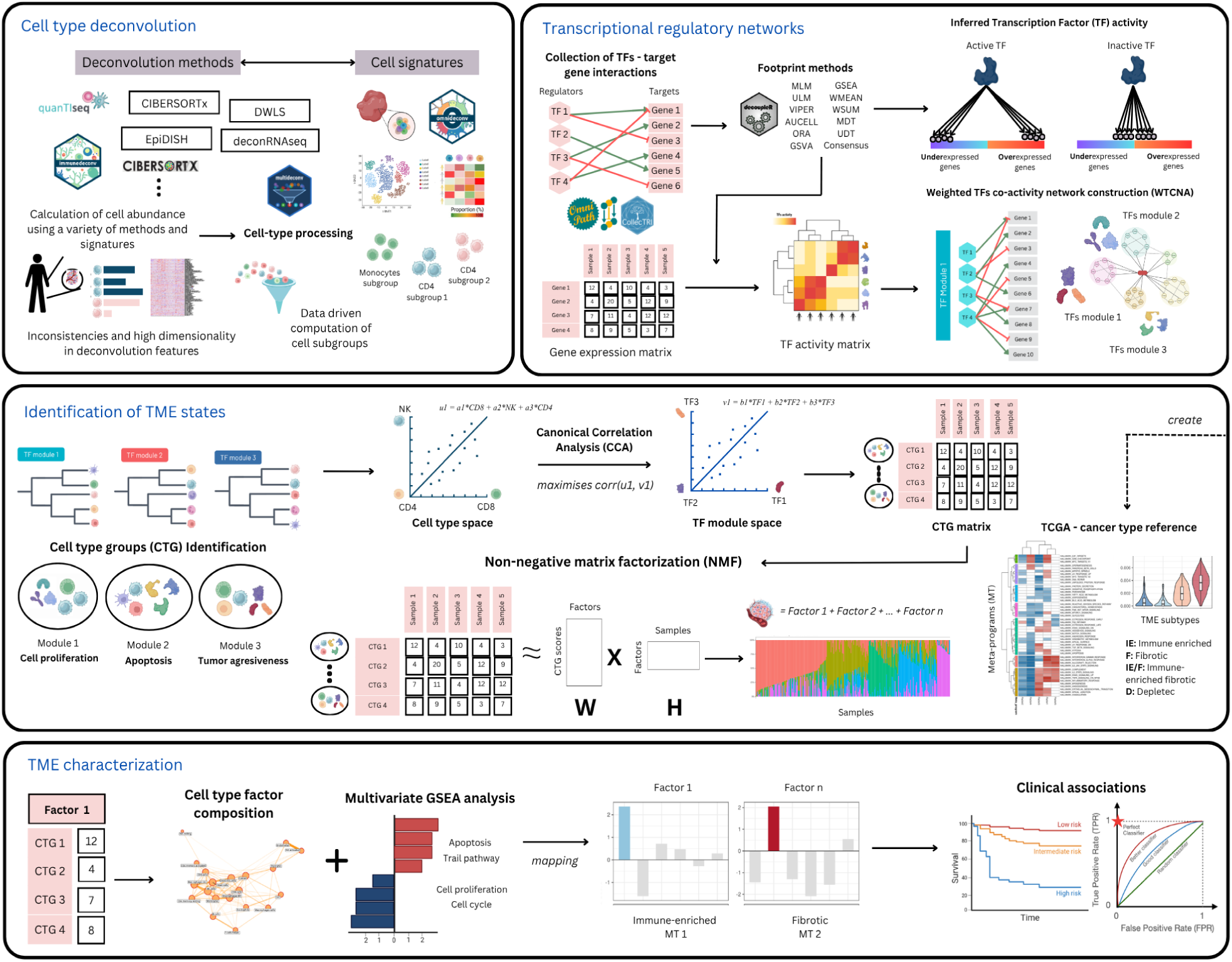
Overview of CellTFusion.

Cell type deconvolution is performed using multideconv by default, an R package that harmonizes outputs from more than six established deconvolution methods and reduces redundancy across correlated cell type estimates through iterative correlation-based filtering (see Methods), yielding a stable, low-redundancy cell-type proportion matrix suitable for downstream integration. CellTFusion is however flexible in its deconvolution input and can accept any pre-computed cell type abundance matrix, allowing users to integrate outputs from alternative deconvolution tools.

In parallel, transcription factor (TF) activity is inferred from normalized gene expression using footprint algorithms coupled with curated gene regulatory networks via OmniPath (see Methods). The resulting TF activity profiles are organized into co-regulatory modules, groups of TFs with correlated activity patterns across samples, using a variation of the weighted gene co-expression network algorithm applied to TF activity (here referred as WTCNA).

Cell type groups are then identified by integrating deconvolution outputs and TF module information. For each TF module, cell types are hierarchically clustered based on their correlation with the module eigengene, and coherent cell-type groupings are identified using dynamic tree cutting (see Methods). The association between each cell group and its corresponding TF module is quantified using canonical correlation analysis (CCA), which identifies the linear combination of cell-type proportions that maximally co-varies with the coordinated TF activity of that module. The resulting composite score for each cell-type group summarizes the coordinated co-variation between cell composition and TF regulatory activity across patients.

Latent factors are derived from the cell type group composite score matrix using non-negative matrix factorization (NMF), compressing the information across all cell groups into a low-dimensional representation in which each factor captures a distinct multicellular regulatory program (see Methods; Supplementary Figure 1).

### Meta-programs derived from TCGA define cancer type specific transcriptional references for interpreting TME latent factors across independent cohorts

To provide a biological reference for the interpretation of CellTFusion latent factors, we applied it to bulk RNA-seq data from 21 TCGA cancer types, building a cancer type specific atlas of what we denote as transcriptional meta-programs (see Methods). PCA of TCGA samples show a large cancer type specific batch effect, which motivated us to perform an independent analysis on each cancer type (Supplementary Figure 2).

Since our primary study cohort in this paper consists of melanoma patients (see below), we illustrate the reference atlas using the SKCM cancer type as an example. CellTFusion identified three latent factors capturing TME heterogeneity across 463 patients, with patients exhibiting mixed factor compositions (Figure 2A). Clustering of Hallmark enrichment results yielded six meta-programs (k=6), each capturing a distinct biologically co-activation pattern (see Methods, Figure 2B). MP1 is defined by broad immune activation, encompassing interferon-γ and interferon-α responses, inflammatory response, complement, IL6-JAK-STAT3 and IL2-STAT5 signaling, TNFα signaling via NFκB, allograft rejection, KRAS signaling, apoptosis, and coagulation. MP2 is characterized by downregulated KRAS signaling, together with early and late estrogen response programs. MP3 captures metabolic and growth signaling activity, including mTORC1 and PI3K-AKT-mTOR signaling alongside adipogenesis. MP4 groups cell cycle and biosynthetic programs, comprising oxidative phosphorylation, MYC targets, E2F targets, G2M checkpoint, and heme metabolism. MP5 reflects proteostatic and metabolic stress programs, including MYC targets variant 2, protein secretion, DNA repair, unfolded protein response, fatty acid metabolism, and cholesterol homeostasis. MP6 defines a stromal and tissue-remodeling program encompassing epithelial-mesenchymal transition, TGF-β signaling, hypoxia, angiogenesis, apical junction and surface remodeling, xenobiotic metabolism, p53 pathway, and myogenesis.

**Figure 2.**
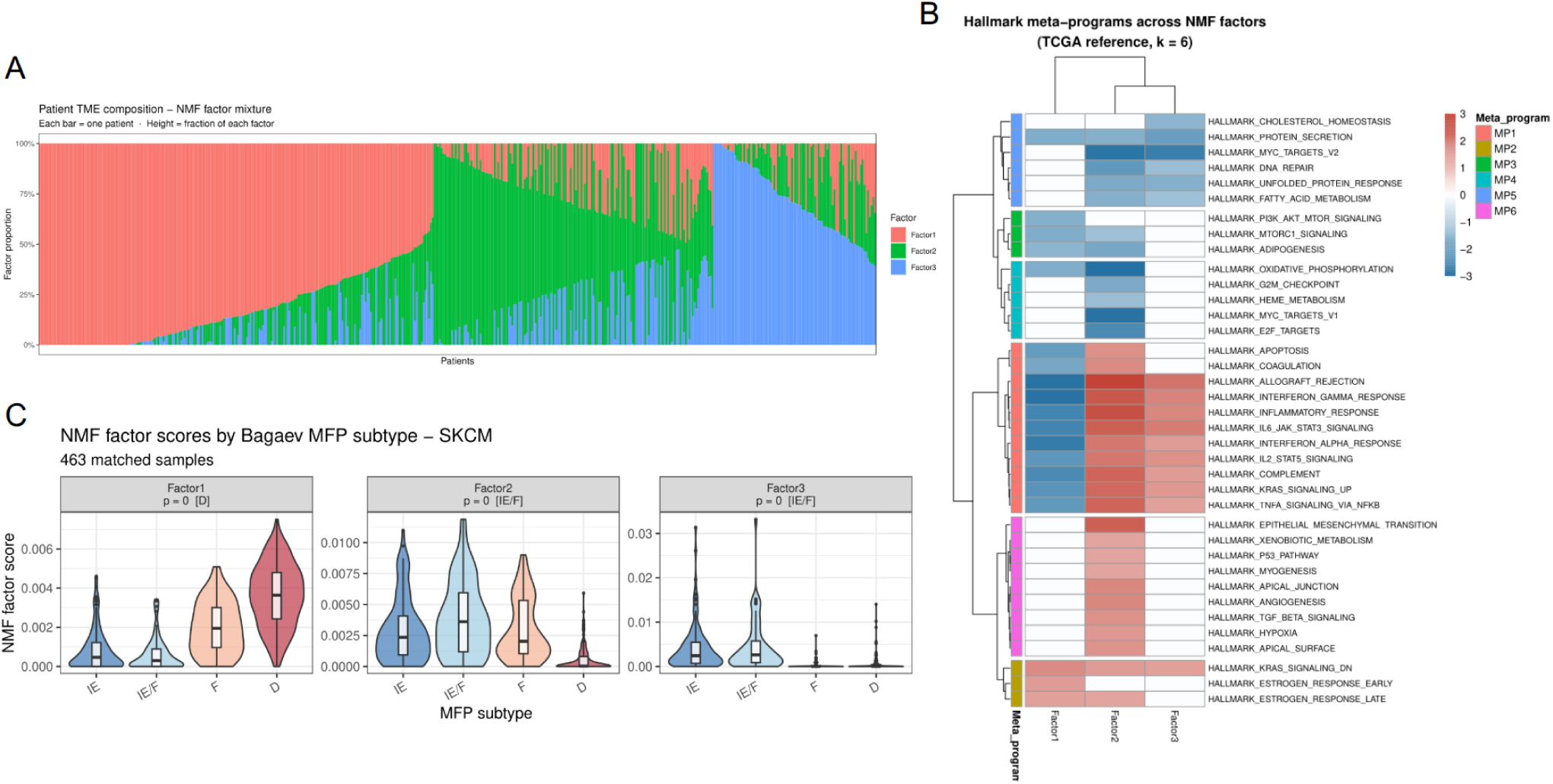
A TCGA-derived transcriptional meta-program reference enables biologically grounded TME characterization across cancer types. **(A)** Patient TME composition barplot for SKCM (n=463). Each bar represents one patient, colored by NMF factor proportion (Factor1, Factor2, Factor3). Patients are ordered by dominant factors. **(B)** Hallmark meta-program heatmap for SKCM (k=6). Rows represent MSigDB Hallmark gene sets, columns represent NMF factors, and color represents the normalized enrichment score (NES). Hallmarks are grouped into six meta-programs (MP1–MP6). **(C)** NMF factor scores stratified by Bagaev et al., 2021 into Molecular Functional Phenotype (MFP) subtype. Violin plots show factor score distributions across IE, IE/F, F, and D subtype groups. Kruskal-Wallis p-values are indicated.

Further characterization of these meta-programs was done using a public TME nomenclature denoted in Bagaev et al., 2021 as Molecular Functional Phenotypes (MFP) and available for TCGA patients (see Methods). All three SKCM factors showed a significant MFP stratification (Krustal-Wallis, p < 0.001; Figure 2C), with Factor 1 most strongly associated with the desert subtype (D) and Factor 2 and 3 with the immune-enriched fibrotic subtype (IE/F). Meta-programs were finally annotated with these TME subtypes using a weighted majority vote approach across contributing factors (see Methods), producing a cancer-type reference for interpreting SKCM CellTFusion factors on independent cohorts.

### CellTFusion identifies recurrent immune and stromal TME programs as transferable predictors of immunotherapy response across six melanoma cohorts

We applied CellTFusion to six public melanoma cohorts treated with immunotherapy (Auslander et al., 2018; Gide et al., 2019; Hugo et al., 2016; Liu et al., 2019; Riaz et al., 2017; Van Allen et al., 2015). PCA analysis revealed strong batch effects between the different cohorts (Supplementary Figure 3), motivating the use of batch aware cell group scoring in CellTFusion (see Methods). CellTFusion identified 5 latent factors capturing the dominant axes of the TME heterogeneity across the combined cohort (Figure 3A).

**Figure 3.**
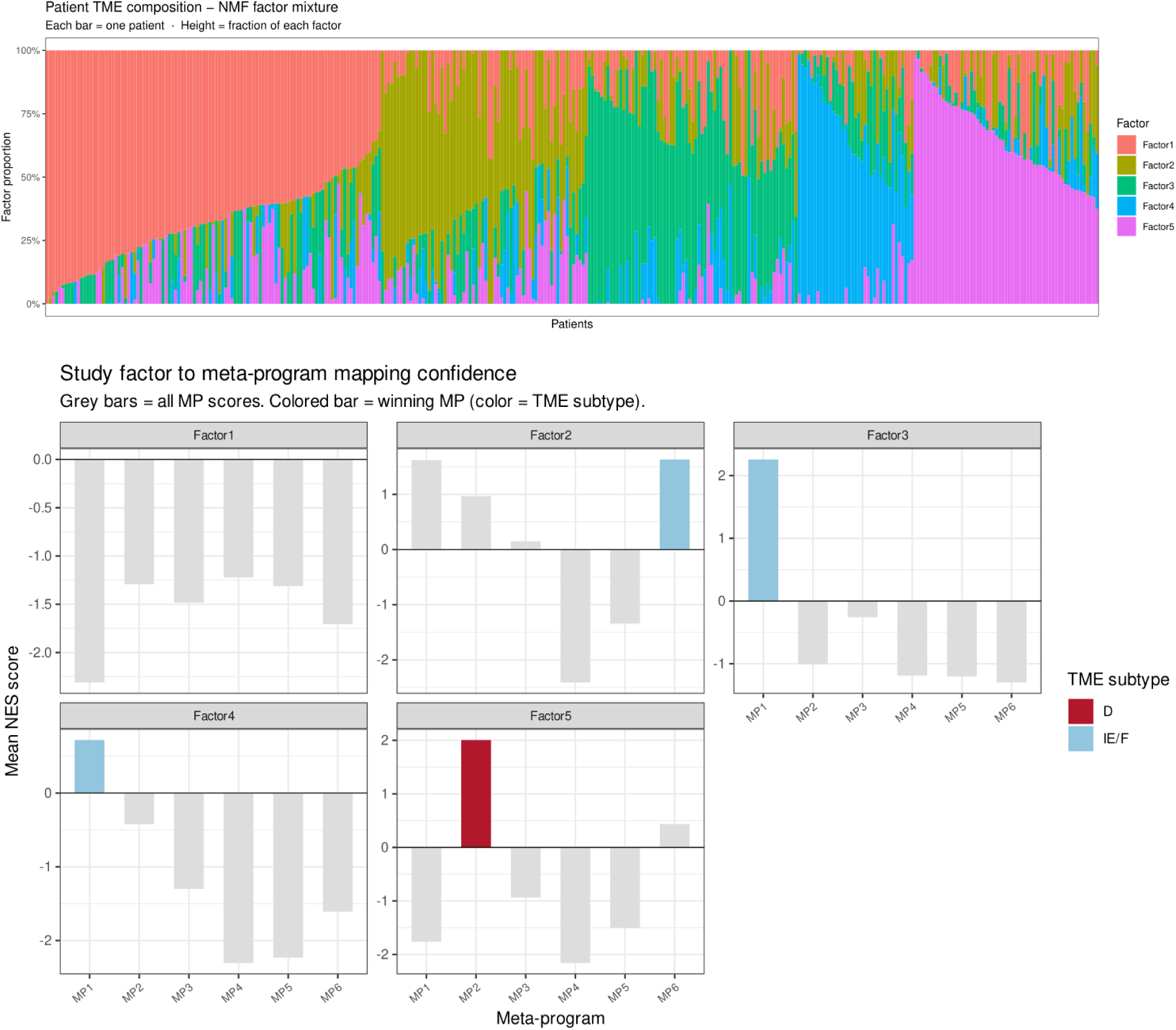
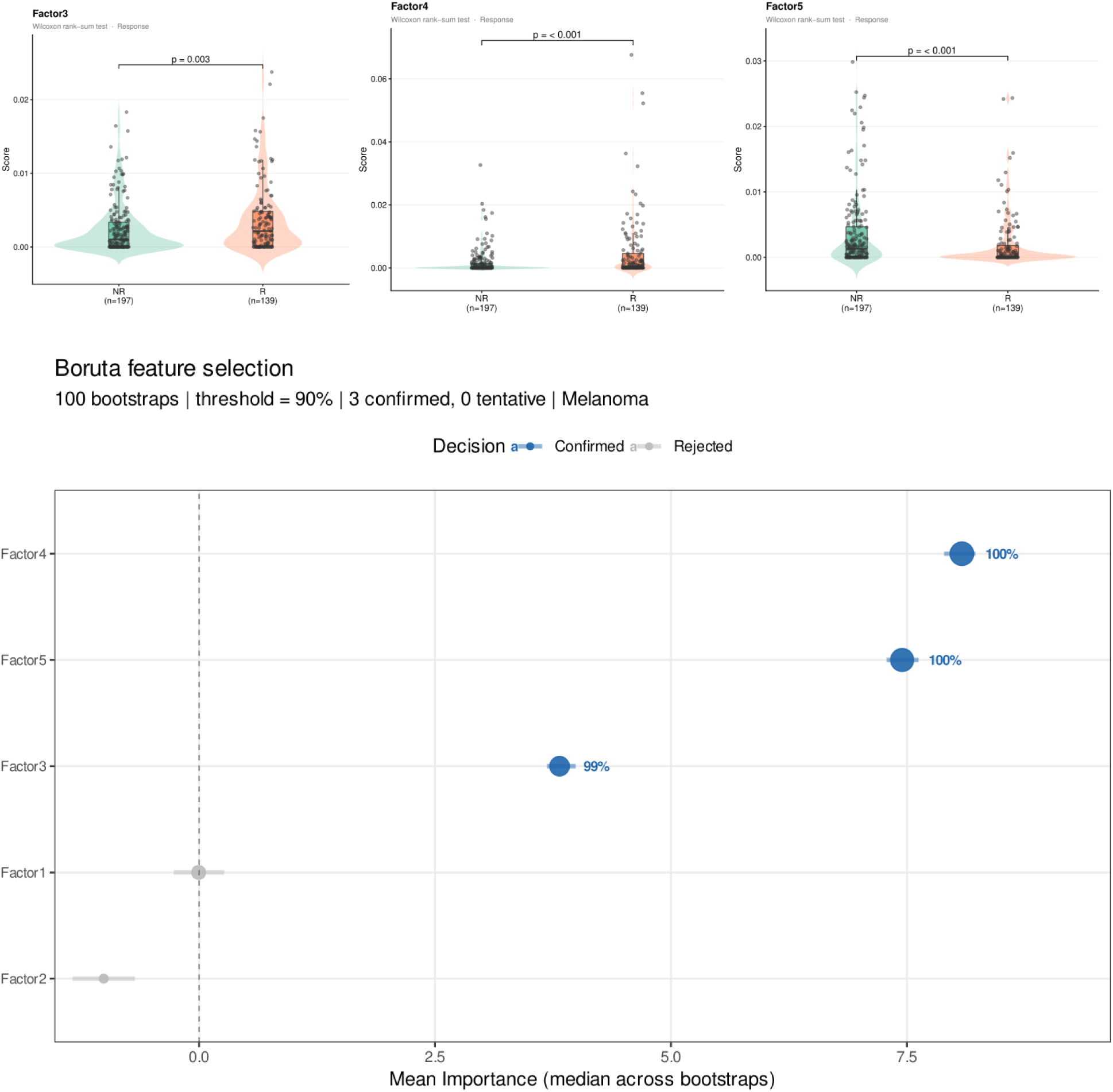
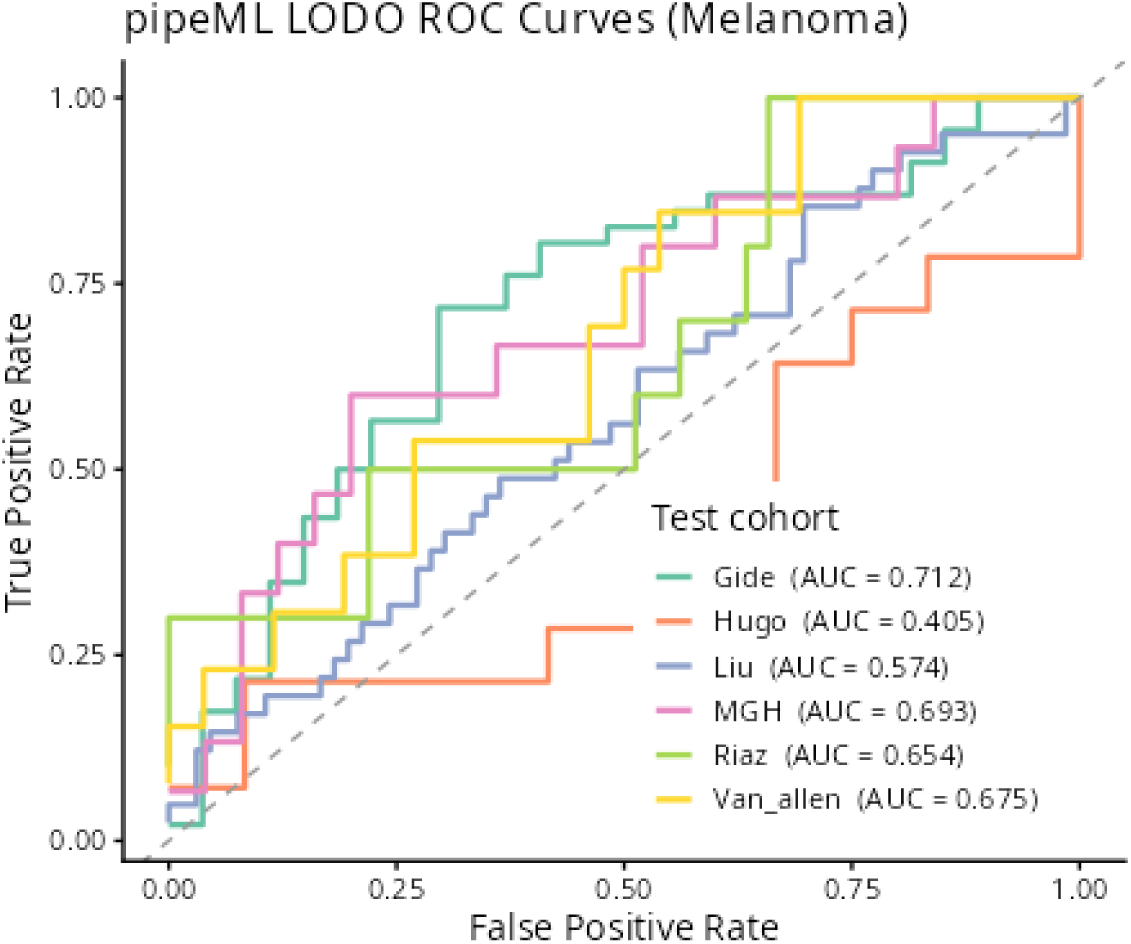
CellTFusion identifies latent TME factors associated with immunotherapy response across six integrated melanoma cohorts. **(A)** Patient TME composition barplot for the combined melanoma cohort. Each bar represents one patient, colored by NMF factor proportion (Factor1–5). Patients are ordered by dominant factors. **(B)** TME factors to SKCM meta-program mapping. For each factor, bars show the mean NES score against each of the six SKCM meta-programs (MP1–MP6). The winning meta-program is colored by its associated TME subtype (blue: IE/F, red: D, grey: uncharacterized). **(C)** NMF factor scores for Factor 3, Factor 4, and Factor 5 stratified by immunotherapy response (NR: n=197, R: n=139). Wilcoxon rank-sum test p-values are indicated. **(D)** Boruta feature selection results across 100 bootstraps (threshold = 90%). Mean importance (median across bootstraps) is shown for each factor; confirmed features are shown in blue, rejected in grey. **(E)** LODO ROC curves for immunotherapy response prediction across five held-out melanoma cohorts using pipeML. AUC values per held-out cohort are indicated in the legend: Gide (0.712), MGH (0.693), Van_allen (0.675), Riaz (0.654), Liu (0.574) and Hugo (0.405).

Biological annotation of these factors was performed using Hallmark NES profiles from GSEA (see Methods) and mapped onto the TCGA-SKCM meta-program reference described above. Factors 2, 3, and 4 were assigned to the IE/F subtype, Factor 5 was assigned to the D subtype, and Factor 1 remained uncharacterized, since no significant association was found to any meta-program (Figure 3B).

Association with immunotherapy response revealed Factors 3, 4, and 5 as significantly discriminating between responders and non-responders (Wilcoxon rank-sum test, p = 0.003, p < 0.001, and p < 0.001 respectively; Figure 3C). Factors 3 and 4 showed higher scores in responders, consistent with their IE/F annotation reflecting active immune programs, while Factor 5 showed higher scores in non-responders, consistent with its D subtype annotation, reflecting an immune-excluded TME. Boruta feature selection confirmed Factor 4 (100%), Factor 5 (100%), and Factor 3 (99%) as robustly relevant predictors across 100 bootstraps (threshold = 90%), while Factors 1 and 2 were rejected (see Methods; Figure 3D).

To assess the predictive value of these factors for immunotherapy response and their robustness, we employed a LODO cross-cohort strategy rather than training and evaluating on the full combined cohort. The latter could lead to overfitting, since CellTFusion factors derived from the complete dataset would already encode information from all samples, making any subsequent within-cohort evaluation optimistically biased. To avoid this, we used pipeML (Hurtado & Pancaldi, 2026), a general pipeline to train prediction models on global features (see Methods). This approach integrates CellTFusion factor computation within each cross-validation fold, ensuring that the held-out cohort does not contribute any information to factor derivation and thus provides an unbiased estimate of cross-cohort performance. As expected, we achieved consistent predictive performance across four out of six held-out cohorts, with AUCs of 0.712 (Gide), 0.693 (MGH), 0.675 (Van_allen), 0.654 (Riaz), 0.574 (Liu) and 0.405 (Hugo) (Figure 3E), demonstrating that these TME factors capture transferable biological signals predictive of immunotherapy response across independent melanoma datasets, with the exception of the Hugo cohort, which showed performance below random expectation.

To evaluate which factors drove predictions in each held-out cohort, we computed SHAP values for each LODO fold model. Since CellTFusion is independently re-run for each training partition under the LODO framework, factors are re-derived for each fold and are not directly comparable across held-out cohorts by their index. Supplementary Figure 4 shows that in the Gide cohort the dominant predictors were Factor 6 and Factor 4, both mapping to the IE/F MP1 program, with high scores of both factors associated with increased likelihood of response. Conversely, Factor 5, mapping to the IE/F MP6 program, and Factor 7, mapping to the D MP2 program, pushed predictions toward non-response. In the MGH cohort, Factor 2 (uncharacterized) and Factor 6 (IE/F, MP6) were the dominant contributors, with high Factor 2 scores associated with non-response and high Factor 6 scores with response. Factor 5 (IE/F, MP1) favored response while Factor 7 (D, MP2) favored non-response. In the Riaz cohort, Factor 1 (uncharacterized) pushed toward non-response, while Factor 7 (IE/F, MP1) pushed toward responders. Factor 5 (IE/F, MP1) also favored response, while Factor 6 (D, MP2) favored non-response, and Factor 4 (IE/F, MP6) also pushed toward non-response. In the Van_allen cohort, Factor 7 (IE/F, MP1) was the dominant predictor favoring response, followed by Factor 6 (IE/F, MP1) also favoring response, while Factor 5 (D, MP2) pushed toward non-response.

Notably, a pattern emerged across cohorts: factors mapping to the IFN-γ/inflammatory IE/F program (MP1) were consistently associated with response, factors mapping to the EMT/TGF-β IE/F program (MP6) showed variable directionality depending on cohort context, and factors mapping to the immune-desert program (MP2) were consistently associated with non-response, suggesting that despite cohort-specific factor differences, CellTFusion coherently identifies the same underlying TME programs as determinants of immunotherapy response across independent melanoma datasets.

### CellTFusion identifies complementary proliferative, and stromal TME programs predictive of immunotherapy response in bladder cancer

We applied CellTFusion to a public bladder cancer cohort (Mariathasan et al., 2018) to assess the generalizability of the CellTFusion framework to other cancer types (Progressive disease PD = 167, Complete response CR = 25). CellTFusion identified 6 factors in this cohort, which were all assigned to the immune-enriched (IE) subtype (Figure 4A) by mapping onto the TCGA-BLCA meta-program reference (see Supplementary Figure 5), reflecting a predominantly immune-active TME in this cohort. The factors captured distinct IE sub-programs: Factors 3, 4, and 6 mapped to MP1 (IFN-γ and IFN-α response, allograft rejection, IL6-JAK-STAT3 signaling), Factor 2 mapped to MP2 (TNFα-NFκB signaling, inflammatory response, epithelial-mesenchymal transition, angiogenesis), and Factors 1 and 5 mapped to MP3 (G2M checkpoint, E2F targets, MYC targets), suggesting a proliferative program within an immune-active context.

**Figure 4.**
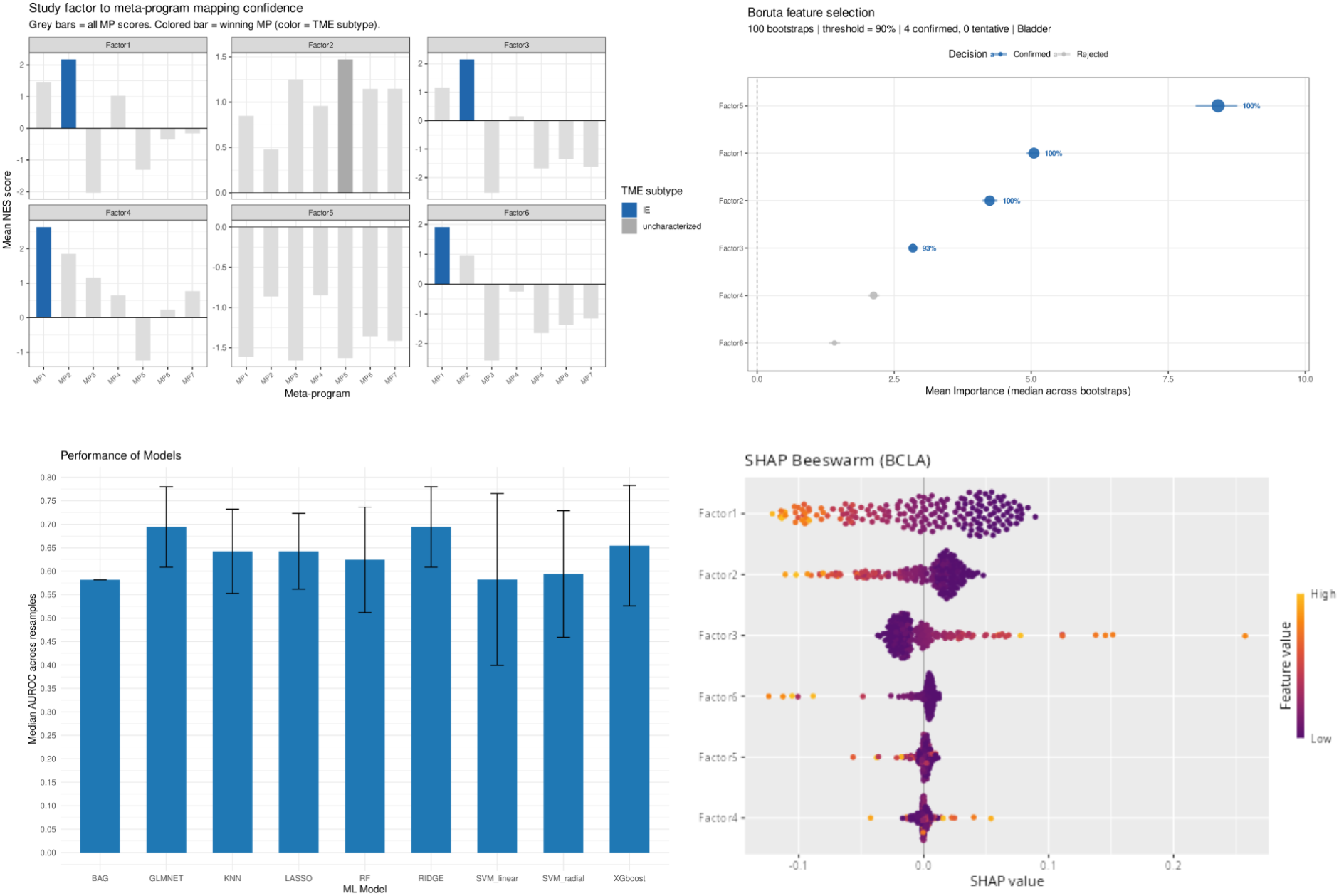
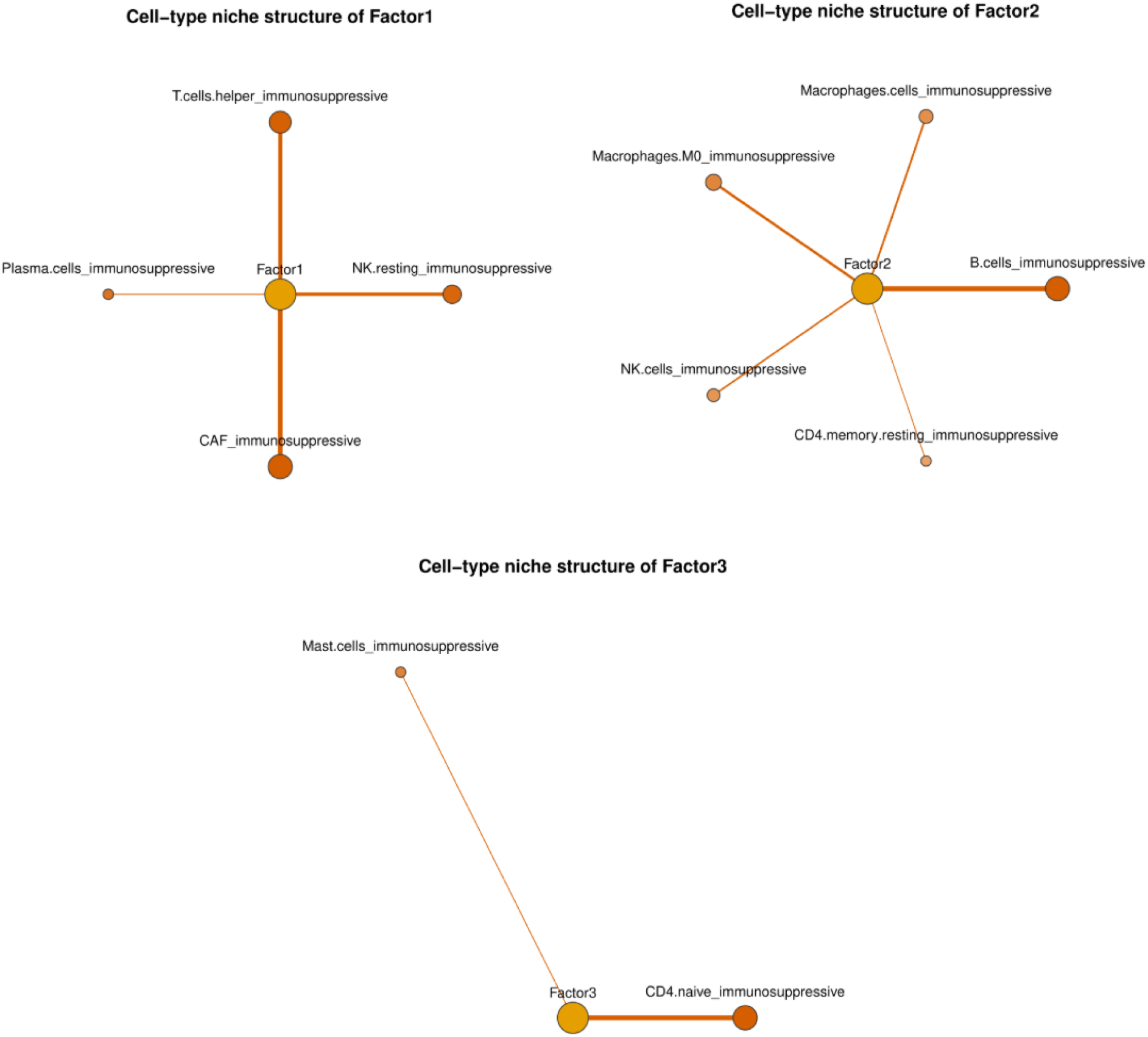
CellTFusion identifies immune-enriched TME factors associated with immunotherapy response in a bladder cancer cohort. **(A)** Study factor to BLCA meta-program mapping. For each factor, bars show the mean NES score against each meta-program (MP1–MP7). The winning meta-program is colored by its associated TME subtype (blue: IE). **(B)** Boruta feature selection results across 100 bootstraps (threshold = 90%). Mean importance (median across bootstraps) is shown for each factor; confirmed features are shown in blue, rejected in grey. **(C)** Performance of machine learning classifiers for immunotherapy response prediction, shown as median AUROC across resamples with error bars indicating variability. **(D)** Left: SHAP feature importance for the top contributing factors, shown as mean absolute SHAP value. Right: SHAP beeswarm plot showing the direction and magnitude of each factor’s contribution to response prediction; color indicates feature value (orange: high, purple: low). **(E)** Cell-type niche structure for the three main factors (Factor 1–3). Nodes represent cell types and the central factor node; edge width reflects the strength of association between the cell type and the factor (see Methods).

Factor scores did not show statistically significant associations with immunotherapy response after Wilcoxon tests, and no significant survival differences were found when stratifying patients by factor scores (Supplementary Figure 6). However, Boruta feature selection confirmed four factors as relevant predictors across 100 bootstraps (threshold = 90%): Factor 5 and Factor 1 (both 100%, highest importance), Factor 2 (100%), and Factor 3 (93%), suggesting that non-linear multi-factor combinations carry the predictive signal (Figure 4B). Machine learning models trained on all factors under cross-validation achieved a median AUROC of ∼0.70 for the best-performing classifiers (GLMNET and RIDGE), confirming that the predictive signal is distributed across factors, rather than concentrated in any single one (Figure 4C). SHAP-based feature importance identified Factor 1, Factor 2, and Factor 3 as the main contributors to predictions (Figure 4D). Dependence plots revealed that high Factor 3 scores pushed predictions toward complete response, consistent with its IFN-γ-driven cytotoxic program (MP1). Conversely, high Factor 1 and Factor 2 scores pushed predictions toward progressive disease, reflecting a tumor proliferative program (MP3) and an inflammatory stromal/EMT program (MP2) respectively, the latter consistent with TGF-β-driven immune exclusion previously identified as a key resistance mechanism in this cohort (Supplementary Figure 7). Taken together, these results suggest that CellTFusion captures a multi-factor TME structure in which two independent adverse programs, tumor proliferation and stromal exclusion, compete against cytotoxic immune activation to determine response, a signal that only emerges through joint multivariate modeling.

Inspection of the cell type niche structure underlying each factor revealed distinct cellular compositions (see Methods; Figure 4E). Factor 1 comprised a mixed immune-stromal niche including T helper cells, plasma cells, resting NK cells, and cancer-associated fibroblasts (CAF), suggesting lymphoid and stromal co-activity. Factor 2 was dominated by a myeloid-rich niche composed by macrophages, B cells, NK cells, and resting CD4 memory T cells, consistent with an immunosuppressive microenvironment characterized by myeloid infiltration and dysfunctional lymphoid populations. Factor 3 showed the most restricted niche, composed only of mast cells and naive CD4 T cells, suggesting a less complex but potentially immuno-regulatory environment. Notably, these results showed that Factors 1 and 2 are both immunosuppressive and lymphoid dysfunctional, one probably to CAF presence and the second to macrophages, providing a cellular explanation for their association with non-responders.

### CellTFusion captures robust and biologically relevant TME communication programs across NSCLC single cell datasets

To evaluate the biological relevance of the cell type groups identified by CellTFusion, we performed cell–cell communication (CC) analysis using LIANA+ (Dimitrov et al., 2024) on a NSCLC single-cell RNA-seq dataset from Vanderbilt University (Senosain et al., 2023). Pseudo-bulk profiles were generated and processed with CellTFusion to derive cell group co-occurrence networks (see Methods).

Restricting the comparison to common cell types, CellTFusion recovered 32% of LIANA’s total interaction frequency weight, with an enrichment score of 0.69, indicating that the shared edges are not enriched for the highest-frequency ligand–receptor interactions in LIANA+ (Figure 5A). While a subset of interactions was shared between the two networks, the majority of edges were unique to each approach, reflecting different information for each of them (Figure 5B. These results suggest that CellTFusion captures aspects of cellular organization that partially overlap, but are not redundant, with direct ligand–receptor interactions, potentially reflecting broader co-regulatory or spatial co-localization signals not captured by ligand–receptor scoring alone.

**Figure 5.**
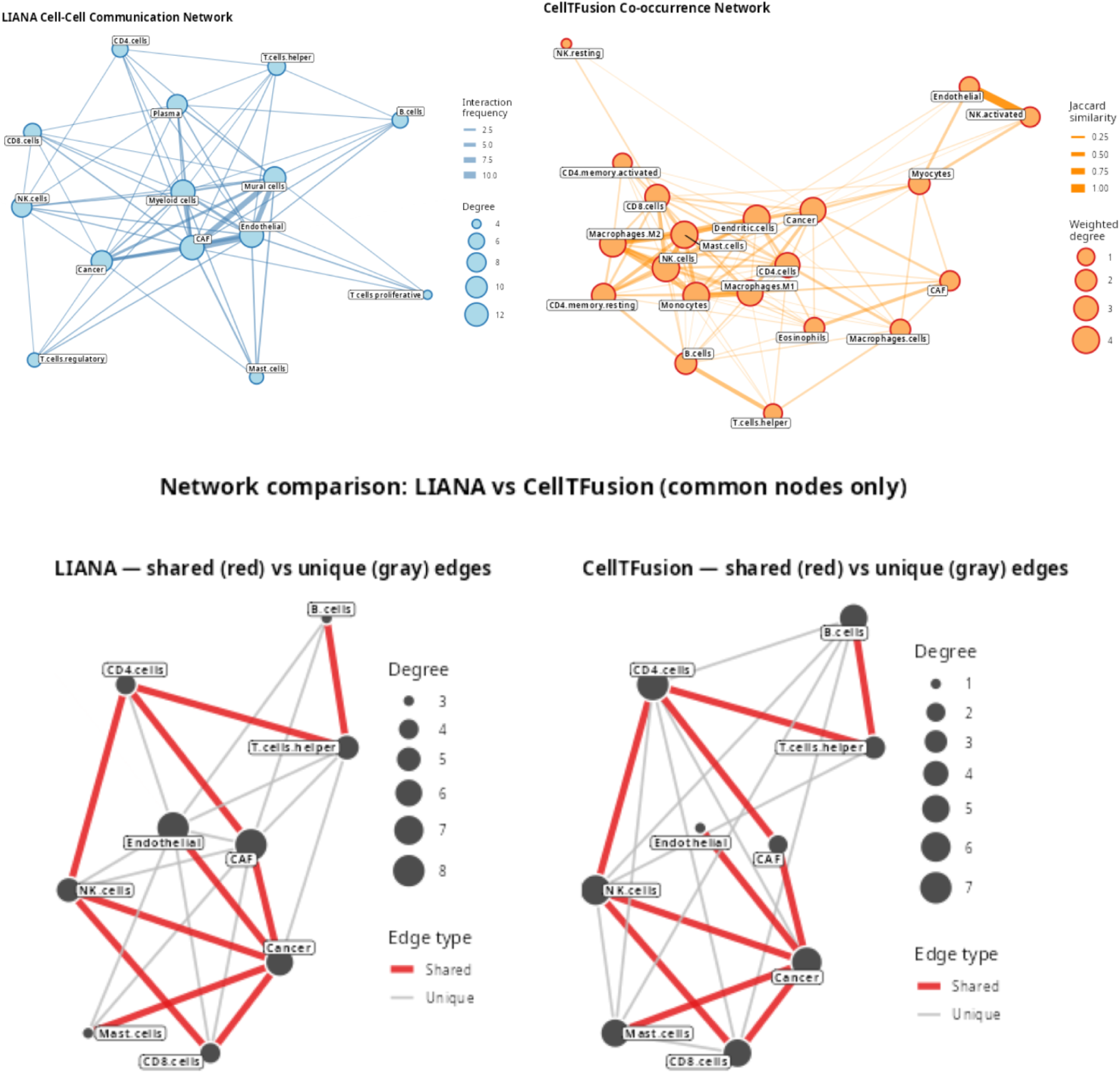
CellTFusion co-occurrence networks partially overlap with but are not redundant to cell–cell communication interactions identified by LIANA+ in the NSCLC scRNA-seq data (Senosain et al., 2023). **(A)** Left: LIANA+ cell–cell communication network inferred from the NSCLC scRNA-seq data. Nodes represent cell types; edge width reflects interaction frequency. Right: CellTFusion co-occurrence network derived from cell group compositions. Nodes represent cell types; edge width reflects Jaccard similarity. **(B)** Network comparison restricted to common cell types. Left: LIANA+ network with shared (red) and unique (grey) edges highlighted. Right: CellTFusion network with shared (red) and unique (grey) edges. Node size reflects degree.

### Benchmarking CellTFusion features against alternative tools to extract insights from the TME reveals transferable profiles for immunotherapy response in a combined melanoma cohort

To benchmark the predictive value of CellTFusion features against alternative approaches for extracting TME information, we used machine learning models to predict immunotherapy response in the combined melanoma cohort using the same LODO framework and CellTFusion latent factors derived above. We compared CellTFusion latent factors against seven alternative feature frameworks: individual TF activity scores, deconvolution estimates, pathway activity scores, and their combinations (TFs + Deconvolution, TFs + Pathways, TFs + Pathways + Deconvolution), cell deconvolution subgroups from multideconv (Hurtado et al., 2025), EaSIeR (Lapuente-Santana et al., 2021), a previously published framework integrating multiple RNA-seq-derived TME views for immune checkpoint blockade response prediction (see Methods) and three variants of the iHet immune heterogeneity score (Lapuente-Santana et al., 2024), a MOFA-derived multimodal signature of TME heterogeneity applied without retraining on the target cohorts (iHet, iHet_excl, iHet_rev; see Methods).

Within-cohort CV resampling analysis showed that most frameworks achieved broadly comparable AUROC distributions, with CellTFusion (median = 0.62) performing similarly to the majority of alternatives. CellTFusion significantly outperformed TF activity scores alone (p < 0.05) and pathway scores alone (p < 0.001), while differences against all combined feature frameworks and deconvolution estimates were non-significant (Supplementary Figure 8). Under the LODO evaluation, CellTFusion achieved the highest median AUROC across held-out cohorts (median = 0.664), outperforming all alternative frameworks including TFs alone (0.635), TFs + Pathways + Deconvolution (0.635), TFs + Deconvolution (0.619), TFs + Pathways (0.614), Subgroups (0.617), iHet_rev (0.589), EaSIeR (0.578), Deconvolution (0.568), iHet_excl (0.566), Pathways alone (0.559), and iHet (0.532), with the Hugo cohort showing reduced performance across all approaches (Figure 6). Together, these results suggest that while CellTFusion performs comparably to combined feature frameworks within a cohort, its TME states capture more transferable biological signals that generalize better across independent datasets, an advantage that only becomes apparent under cross-cohort evaluation.

**Figure 6.**
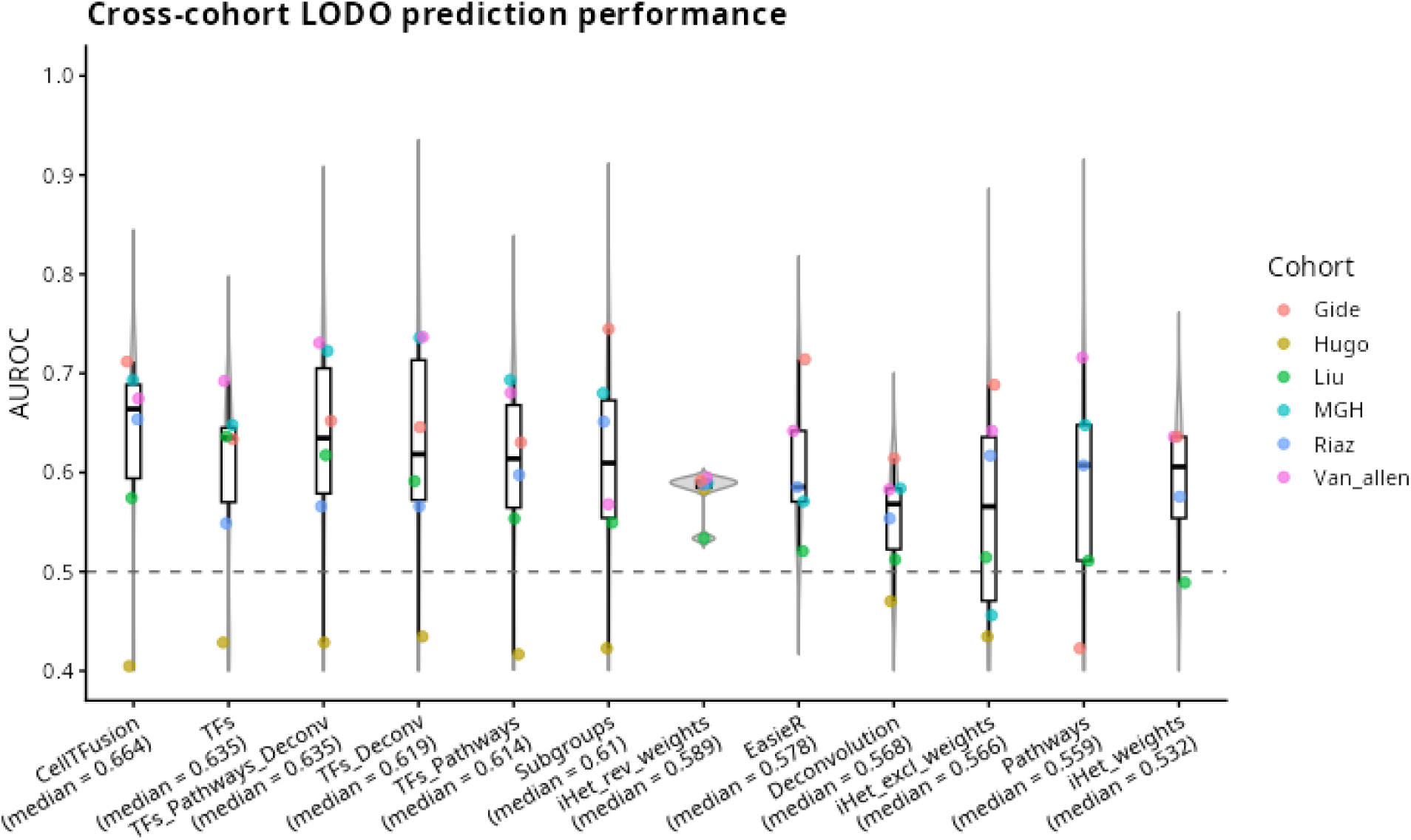
Benchmarking CellTFusion against alternative TME feature frameworks for immunotherapy response prediction in the combined melanoma cohort. Cross-cohort LODO prediction performance. Each violin plot shows the distribution of per-cohort AUROC values across held-out datasets for each feature framework. Colored dots represent individual cohorts (Gide, Hugo, Liu, MGH, Riaz, Van_allen). The dashed line indicates random classifier performance (AUROC = 0.5). Median AUROC across cohorts is indicated for each framework: CellTFusion (0.664), TFs (0.635), TFs + Pathways + Deconvolution (0.635), TFs + Deconvolution (0.619), TFs + Pathways (0.614), Subgroups (0.617), iHet_rev (0.589), EaSIeR (0.578), Deconvolution (0.568), iHet_excl (0.566), Pathways (0.559), and iHet (0.532).

### An interactive RShiny application enables user-friendly exploration and interpretation of CellTFusion results

To facilitate usage, exploration, and interpretation of CellTFusion by non-coding users, we developed an interactive RShiny application. This application provides a guided interface through the main outputs of the pipeline, organized around three levels. At the cell group level, users can explore the composition of each identified cell group, inspect the TF–cell-type association networks underlying each group, and examine the canonical correlation scores across samples. At the latent factor level, users can visualize the NMF factor mixture per patient, inspect top-contributing cell groups per factor, and explore the Hallmark enrichment profiles used for meta-program mapping. At the clinical level, users can assess associations between factor scores and clinical variables including immunotherapy response and survival, and project pre-computed cell group models onto new cohorts without rerunning the full pipeline. Together, the application allows code-free interrogation, interpretation and application of CellTFusion outputs to independent datasets through a point-and-click interface (Figure 7)

**Figure 7.**
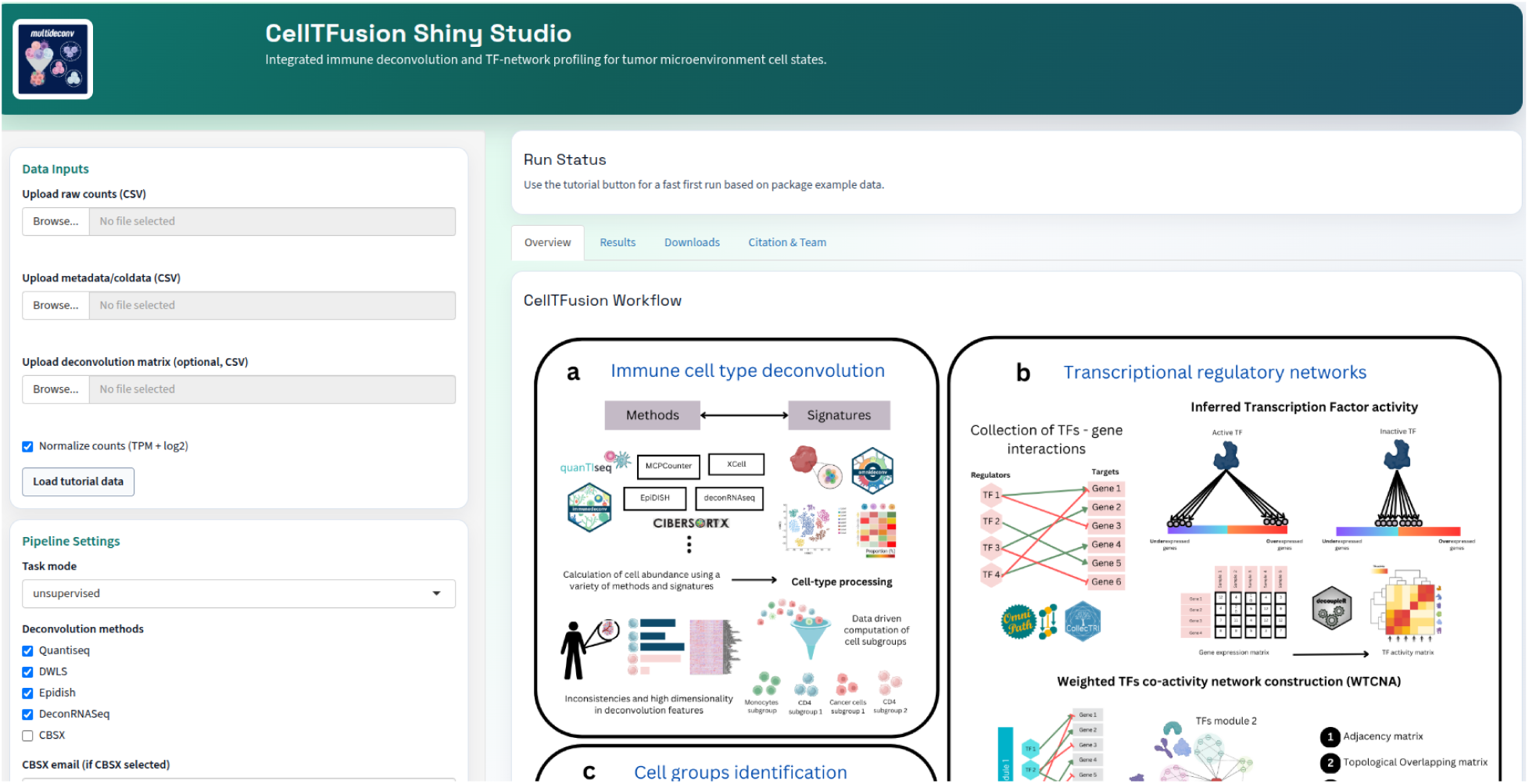
CellTFusion R-shiny application interface. Overview of the CellTFusion interactive RShiny application. The interface provides point-and-click exploration of cell group compositions, TF–cell-type association networks, NMF latent factor structures, and clinical associations across patient cohorts.

## Discussion

The TME has emerged as a central determinant of cancer progression and response to immunotherapy, shifting the focus from cancer cells toward the study of the complex multicellular ecosystem that shapes tumor behavior. Despite this, current computational approaches that aim to characterize the TME from bulk RNA-seq data treat its components in isolation (e.g. cell type deconvolution, pathway activities) rather than capturing the coordinated regulatory programs that define the functional multicellular states (Lapuente-Santana et al., 2021; van Santvoort et al., 2025). Recent work has proposed a new view of the hallmarks of cancer, stating that they should be understood not as cell-intrinsic traits but as emergent ecosystem properties, shaped by the spatial organization and intercellular coordination of the cells present in the tumor niche (Quail & Joyce, 2026; Sibai et al., 2026; Swanton et al., 2024). Under this new framework, characterizing the TME requires methods capable of identifying coordinated programs spanning multicellular environments simultaneously. In this study, we describe CellTFusion, an approach that aims to unravel these functional multicellular states that reflect coordinated biological states from the TME from bulk RNA-seq data.

While recent tools for bulk TME characterization have provided important insights into tumor composition by estimating cell-type abundances, pathway activities or immunoscores; they treat these levels as independent sources of information (Lapuente-Santana et al., 2024; Luca et al., 2021). In contrast, CellTFusion explicitly bridges TF activity with cell-type abundance through canonical correlation analysis, giving a score to the degree to which coordinated TF activity co-varies with cell type composition across patients, and then compressing this joint signal into low-dimensional latent factors via non-negative matrix factorization. The resulting factors capture shared functional states rather than solely cellular co-abundance, providing a representation that is biologically interpretable, statistically robust, and transferable across independent cohorts. Critically, CellTFusion is not a replacement for existing deconvolution tools, but a flexible integration layer that accepts any precomputed cell type abundance matrix as input and adds a regulatory layer via TF activity to any deconvolution strategy the user applies.

Benchmarking against other TME characterization approaches confirmed that CellTFusion scores are not redundant with existing features but provide complementary predictive signals that can improve both performance and interpretability compared to any single layer alone. Importantly, the advantage of CellTFusion over individual feature frameworks was most evident under cross-cohort LODO evaluation, where CellTFusion achieved the highest median AUROC on prediction, despite performing comparably during the training. This suggests that the CellTFusion factors capture more generalizable biological structure than the raw concatenation of individual feature layers, improving transferability across independent datasets.

A major strength of CellTFusion is that it produces a mixture representation of the TME rather than a categorical assignment. Each patient is described as a weighted combination of latent factors, each capturing a distinct coordinated hallmark program. A critical consequence of this mixture representation is that it resolves functionally distinct programs within the same broad TME subtype category. For example, in our melanoma analysis, both the IFN-γ/inflammatory program (MP1) and the EMT/TGF-β stromal program (MP6) were classified as IE/F under the Bagaev categorical nomenclature, yet they showed consistently opposing associations with immunotherapy response across independent cohorts (MP1 favoring response and MP6 favoring non-response). This distinction would have been invisible under a categorical TME subtype assignment, where both programs would have been collapsed into the same IE/F label. The ML model implicitly captures this competition: neither program alone was sufficient for prediction, and the model required simultaneous information about both protective and adverse programs to discriminate responders from non-responders according to our SHAP analysis, consistent with a TME ecosystem where opposing functional states co-exist and their relative balance determines therapeutic outcome. This mixture aims to look at bulk data as an aggregate of multiple co-occurring ecosystem states competing between each other.

Consistent with this, in both melanoma and bladder cohorts our results showed heterogeneous factor compositions where patients co-expressed multiple functional programs simultaneously. In melanoma, IE/F and D factors co-existed within the same patients, with the balance between IFN-γ-driven cytotoxic programs and EMT/TGF-β stromal programs determining response. In bladder cancer, all six factors mapped to the IE subtype yet captured functionally distinct sub-programs, including proliferative, inflammatory-stromal, and cytotoxic; whose joint activity, rather than any single factor, was predictive of response. In both cases, no single factor was sufficient to discriminate responders from non-responders in univariate analysis (Wilcoxon tests), and the predictive signal only emerged through joint multivariate modeling, consistent with the view that response is determined by the relative dominance of competing ecosystem states rather than the presence of any single cell type or pathway.

The biological relevance of the identified cell type groups was further supported through a scRNA-seq validation analysis where we showed how CellTFusion features captured high-confidence ligand-receptor interactions as well as other types of interactions suggesting it can capture mechanisms that go beyond L-R mediated cell-cell communication. Notably, the enrichment score of 0.69 and the partial overlap between CellTFusion and LIANA+ networks indicate that CellTFusion co-occurrence patterns are not simply a proxy for ligand-receptor communication but reflect a broader organizational principle of cellular co-regulation, potentially encompassing spatial co-localization, shared cytokine environments, and transcriptional crosstalk that are not captured by explicit ligand-receptor scoring alone.

Despite these advantages, CellTFusion inherits the intrinsic limitations that are inherent in bulk RNA-seq data. Bulk measurements collapse a mixture of cells into a single averaged profile, making it hard to fully disentangle with confidence the TME. Furthermore, bulk measurements are inherently sensitive to tumor purity, as the dominance of the tumor cell signal can mask or distort the weaker contributions of less abundant cell types in the TME. Nevertheless, bulk RNA-seq remains the most widely available and cost-effective data modality in both research and clinical settings, with thousands of publicly available datasets spanning diverse cancer types and treatment contexts. Compared to single-cell RNA-seq or spatial omics, which offer higher resolution but at substantially greater cost, technical complexity, and limited clinical scalability, bulk RNA-seq still represents a powerful tool for TME characterization at the scale required for biomarker discovery and clinical translation.

A second class of limitations concerns the reliance on prior-knowledge databases. TF activity is inferred from target gene expression using footprint-based methods, which are regulatory proxies whose accuracy relies on the completeness and quality of the underlying gene regulatory network. Similarly, cell type abundance estimates depend on the specificity and coverage of reference signatures. To mitigate these risks, we used multideconv, a tool which combines multiple deconvolution methods and signatures to give subgroups estimates of cell proportions (Hurtado et al., 2025), providing more robust and less biased abundance estimates, and used CollecTRI (Müller-Dott et al., 2023) as the default curated regulon database, one of the most comprehensive and curated collections of signed transcriptional regulatory interactions currently available.

In summary, our results demonstrate that functional multicellular ecosystems can be inferred from bulk RNA-seq data, transferred across independent datasets, and linked to clinically relevant outcomes. CellTFusion provides performance comparable to existing approaches for TME characterization, while offering improved generalisability and complementary biological insight, holding promise for the identification of predictive biomarkers of therapeutic response.

## Methods

### CellTFusion inputs

CellTFusion requires as primary input a matrix of raw gene expression derived from bulk RNA-seq. Raw counts are internally normalized to transcript per million (TPM) using the ADImpute package v1.22.0 (Leote, 2026) and log_2_-transformed prior to transcription factor activity inference, while TPM counts are used as input for deconvolution.

### Cell type deconvolution

Cell type proportion estimates are computed using multideconv v.0.0.1 (Hurtado et al., 2025), an R package that harmonizes and integrates results from multiple bulk RNA-seq deconvolution methods, reducing redundancy across method outputs through iterative correlation-based grouping. Alternatively, users may supply a precomputed deconvolution matrix directly. In both cases, the resulting cell type proportion matrix is used as input for downstream cell group identification.

### Transcription factor activity inference

Transcription factor (TF) activity was inferred from bulk RNA-seq gene expression using the decoupleR package v2.9.7 (Badia-i-Mompel et al., 2022), which was run in multi-method mode combining multivariate linear model (MLM), univariate linear model (ULM), and weighted sum (WSUM), the top-performing methods in the decoupleR benchmark, and computing a consensus activity score across them. We used CollecTRI as the regulatory network, a collection of transcriptional regulatory interactions, which provides regulons containing signed TF - target gene interactions compiled from 12 different resources (Müller-Dott et al., 2023) via OmnipathR v4.1.0 (Türei et al., 2016). Alternatively, users may provide a custom regulatory network, or supply an ARANCE-inferred network (see below) (Lachmann et al., 2016; Margolin et al., 2006). The resulting TF activity matrix was z-score normalized to account for variability in regulon size.

### Cancer-type-specific regulatory networks via ARACNe-AP

As an alternative to curated regulon databases, CellTFusion supports the use of cancer-type-specific gene regulatory networks for the calculation of TF activities, inferred for the TCGA bulk RNA-seq data for the 21 cancer types using ARACNe-AP, an implementation of the Algorithm for the Reconstruction of Accurate Cellular Networks (ARACNe) based on an Adaptive Partitioning strategy for estimating Mutual Information (MI) between gene pairs. Unlike curated databases that aggregate interactions across diverse experimental contexts, ARACNe derives regulons directly from the transcriptional covariation structure of a given cancer cohort, producing networks that reflect the regulatory landscape of that specific tumor type (Lachmann et al., 2016; Margolin et al., 2006). For each TCGA cancer type, raw counts were downloaded using the TCGAbiolinks R package v2.40.0 (Colaprico et al., 2016) and further normalized to log_2_(TPM+1) to be used as input expression matrices, with the candidate regulator list restricted to CollecTRI TFs. Networks were inferred using default values from the pipeline, corresponding to 100 bootstrap iterations at a mutual information significance threshold of p < 1×10⁻⁸, followed by consolidation into a final consensus network. Since ARACNe-AP outputs are unsigned, the mode of regulation for each TF–target edge was inferred from the sign of the Spearman correlation between regulator and target expression across samples, with edges of undefined or zero correlation discarded. The resulting signed networks, pre-computed for each supported TCGA cancer type, are bundled within the CellTFusion package and loaded automatically when the user sets *TF.collection = “ARACNE”* alongside the corresponding *cancer_type* argument, when computing TF activity.

### Weighted TFs co-activity networks (WTCNA)

Transcriptional regulation is governed by complex, highly interconnected networks of TFs that cooperatively control gene expression programs and cellular phenotypes. Beyond regulating target genes, TFs frequently regulate each other, forming higher-order regulatory circuits whose coordinated activity underlies biological processes. To identify such coordinated regulatory programs, we adapted the weighted gene co-expression network analysis (WGCNA) (Langfelder & Horvath, 2008) to inferred TF activity subnetworks, resulting in weighted transcription factor co-activity network analysis (WTCNA) using the R package WGCNA v1.74. In this framework, pairwise correlations between TF activity scores derived from RNA-seq data are used to construct a weighted network, which is subsequently partitioned into modules of highly co-active TFs. Each module is summarized by its eigengene, representing the first principal component of TF activities within the module, and serves as a low-dimensional representation of coordinated TF regulation suitable for downstream association analyses with phenotypic traits.

To account for cohort-specific effects and enhance robustness across datasets, WTCNA was performed either on individual cohorts or using a consensus network approach when multiple cohorts were available. In the consensus setting, cohort-specific soft-thresholding powers were first selected to approximate scale-free topology independently within each cohort. TF activity matrices were then combined using WGCNA’s blockwise consensus module detection framework to identify TF modules conserved across cohorts. Module eigengenes were computed separately for each cohort, scaled within cohorts to remove batch-driven differences in magnitude, and subsequently concatenated across cohorts for downstream analyses. This strategy enables the identification of reproducible TF co-activity modules while mitigating technical and cohort-specific confounding effects, yielding biologically interpretable and comparable regulatory features across heterogeneous datasets.

### Cell type groups identification

Cell type group associations were identified by computing module–trait relationships between TF module eigengenes and cell-type abundance estimates derived from bulk RNA-seq deconvolution. By default, pairwise associations were quantified using pearson correlation coefficients; when specified, batch effects (e.g. cohort of origin) were explicitly controlled for by computing partial correlations with the batch variable included as a covariate. For each module–cell-type pair, statistical significance was assessed using Student’s t-distribution for standard correlations or asymptotic p-values for partial correlations.

### Cell type groups composite score

For each TF module, cell type groups were defined by hierarchically clustering the correlation matrix between the TF module eigengenes and the cell type abundance estimates. The resulting dendrogram was automatically cut using the *cutreeDynamic* algorithm from the dynamicTreeCut v1.63-1 package (Langfelder & Horvath, 2008), which adaptively determines the number and boundaries of clusters based on the local shape of the dendrogram rather than a fixed distance threshold, using a tree-based method with default parameters (deep split parameter = 4, minimum cluster size = 3). To quantify the strength and direction of association between each cell type group and its corresponding TF module, we employed canonical correlation analysis (CCA). TF module eigengenes were first used to determine the global direction of association with the cell group. When a negative association was detected, the cell group matrix was inverted prior to CCA, such that the resulting canonical weights implicitly encoded the correct directionality.

CCA was then applied between the scaled cell group abundances and the activity profiles of all TFs belonging to the corresponding module, identifying the linear combination of cell types that maximized the correlation with the coordinated TF activity. Associations with weak canonical correlations or, more specifically, those that did not reach statistical significance were discarded. Statistical significance was calculated based on a permutation test (999 permutations, p > 0.05; 999 chosen to yield a minimum p-value of 0.001). The resulting first canonical variate of the cell group was retained as a composite score summarizing the coordinated association between cell composition and TF regulatory activity.

To control for potential cohort- or batch-specific confounding effects, batch correction was performed when indicated by regressing out the batch variable from cell group abundances, TF module eigengenes, and individual TF activity profiles prior to CCA. This ensured that composite scores reflected biological covariation rather than technical or cohort-driven effects.

### Latent factor inference

To identify reduced representations that can summarize the information contained on the cell groups scores, we applied Non-negative-Matrix Factorization (NMF) using the RcppML package v0.3.7.1 (DeBruine et al., 2024). The input consisted of the cell groups matrix (samples x cell groups), where each entry represents the CCA composite score of a given cell group in a sample. Since NMF requires non-negative inputs, each cell group score was decomposed into its positive and negative direction components. These two components were concatenated column-wise, yielding a non-negative input matrix of samples x (2 x cell groups).

After NMF, the final matrix was decomposed as A ≈ W × diag(d) × H, where W contains feature weights (cell groups × rank) and H contains sample-level factor scores (rank × samples). Rank corresponds to the number of factors that should be inferred by the algorithm. If not supplied by the user, this is estimated automatically by fitting NMF models across ranks 2 to 8 and selecting the rank at the elbow of the reconstruction mean squared error (MSE) curve, identified as the point of minimum second-order difference. The resulting factor score matrix Z (samples × rank) and weight matrix W (features × rank) were retained for downstream characterization.

### CellTFusion niches identification

To characterize the cellular composition of each factor, we identified cell type niches by mapping top-contributing cell type groups to their underlying cell composition and assessing enrichment scores relative to the background frequencies. For each factor, top-contributing cell type groups were selected using a quantile threshold applied to the NMF weight matrix W. Each cell type group was then mapped to a binary composition matrix indicating which cell types it contains. Edge weights connecting a factor to each cell type were computed as the cumulative sum of NMF weights across all top cell groups containing that cell type, reflecting the overall strength of association between the factor and each cell type. To retain only biologically relevant associations, cell types were kept only if their frequency among the top contributing cell groups exceeded their background frequency by a default factor of 1.5.

### TCGA TME meta-program reference

We constructed a reference set of TME meta-programs from the The Cancer Genome Atlas (TCGA) bulk RNA-seq data for 21 cancer types (bladder urothelial carcinoma [BLCA], breast invasive carcinoma [BRCA], cervical squamous cell carcinoma [CESC], cholangiocarcinoma [CHOL], colon adenocarcinoma [COAD], esophageal carcinoma [ESCA], head and neck squamous cell carcinoma [HNSC], kidney chromophobe [KICH], kidney renal clear cell carcinoma [KIRC], kidney renal papillary cell carcinoma [KIRP], pan-kidney [kidney], liver hepatocellular carcinoma [LIHC], lung adenocarcinoma [LUAD], lung squamous cell carcinoma [LUSC], pan-lung [LUNG], prostate adenocarcinoma [PRAD], rectum adenocarcinoma [READ], skin cutaneous melanoma [SKCM], stomach adenocarcinoma [STAD], thyroid carcinoma [THCA], and uterine corpus endometrial carcinoma [UCEC]). Raw counts were downloaded and preprocessed as described above, and CellTFusion was applied to each TCGA cancer cohort to derive NMF latent factors and their sample-level scores. Each factor was characterized by fitting a multivariate linear model to the log_2_(TPM + 1) expression matrix using all factor scores simultaneously as covariates, using the lmFit and eBayes functions from the R package limma v3.62.2 (Ritchie et al., 2015), and performing preranked GSEA against the MSigDB Hallmark collection (50 gene sets) (Liberzon et al., 2015) extracted via the R package msigdbr v26.1.0 and using the t-statistics as the gene ranking. This produced a Hallmarks x factors normalized enrichment score (NES) matrix, in which non-significant (p value > 0.05) Hallmarks were set to zero.

Meta-programs were derived by clustering Hallmark gene sets across each TCGA factor. Pairwise Euclidean distances between Hallmark NES profiles were computed and hierarchical clustering was performed using Ward.D2 linkage using the R package stats v4.4.1. The number of meta-programs k was determined automatically by the elbow method, which consists of identifying the elbow of the within-cluster sum of squares (WSS) curve via the minimum of its second derivative. Each resulting cluster defines a meta-program as a set of Hallmark gene sets with recurrently co-occurring enrichment profiles across TCGA factors, capturing dominant biological processes categories across the TCGA cancer samples.

We then used a previously defined classification of the TME (referred originally as Molecular Functional Phenotypes (MFP)) to assign patients into different subtypes: Immune-Enriched Fibrotic (IE/F), Immune-Enriched Non-Fibrotic (IE), Fibrotic (F) and Desert (D) (Bagaev et al., 2021). The MFP subtype associated with each TCGA patient was provided by the original work. For each NMF factor, a Kruskal-Wallis test was performed to assess whether factor scores differed significantly across the four MFP groups. Factors with a significant test (p < 0.05) were assigned the MFP subtype corresponding to the group with the highest median factor score, reflecting which TME subtype is most strongly associated with high activity of that factor. Factors that did not reach significance were labeled as *uncharacterized*.

Meta-programs were subsequently annotated with a TME subtype via a weighted majority vote procedure. Each meta-program receives contributions from all TCGA factors that map to it, where the mapping score of each factor to that meta-program, defined as the mean NES of the meta-program’s constituent Hallmarks in that factor, serves as its weight. Only factors with a positive mapping score contribute to the vote. For each MFP subtype, the weights of all contributing factors carrying that subtype label are summed, and the subtype with the highest cumulative weight is assigned to the meta-program. Meta-programs for which the winning label is *uncharacterized*, or for which no factors contribute a positive mapping score, are also labeled *uncharacterized*. The resulting reference comprises the Hallmark composition of each meta-program and its associated TME subtype for further use as a reference for the characterization of NMF factors derived from independent study cohorts.

### TME states characterization

For the biological characterization of NMF latent factors, we applied the same multivariate limma-GSEA framework described above, producing a Hallmarks × factors NES matrix covering all 50 MSigDB Hallmark gene sets. Each study factor was then mapped to the TCGA meta-program reference (see above) by computing, for each meta-program, the mean NES of its constituent Hallmark gene sets in that factor. The meta-program with the highest mean NES was assigned as the primary match. The TME subtype annotation was inherited directly from the matched reference meta-program, providing a biologically grounded and TCGA-anchored interpretation of each study factor, without requiring TCGA samples to be present in the study cohort.

### Cell type group and latent factor projection

To map cell groups and latent factors identified in a study cohort onto another independent cohort, we perform a projection in two sequential steps. First, cell group scores are computed for the test cohort by applying the canonical weights estimated during training to the corresponding deconvolution features of the new samples. Specifically, for each cell group, the relevant deconvolution features are extracted from the test data, scaled using the mean and standard deviation from the training cohort, and multiplied by the first canonical coefficients from the original CCA. Features absent in the test cohort or with zero variance are excluded, and only features shared between training and test are used. This produces a cell group score matrix for the test samples that is directly comparable to the training scores.

Second, test cell group scores are projected onto the trained NMF latent space using the Moore-Penrose pseudoinverse of the basis matrix W. Since RcppML stores the NMF decomposition as A ≈ W × diag(d) × H, the diagonal scaling vector d is first absorbed into W before computing the pseudoinverse, ensuring that projected scores are on the correct scale. The projection is restricted to features shared between training and test. The result is a matrix of latent factor scores for the test samples directly comparable to those of the training cohort.

### Supervised and unsupervised cell group analysis

CellTFusion operates in two modes depending on whether a clinical contrast is provided. In the unsupervised mode, TF activity is computed across all samples and TFs are further clustered into WTCNA modules (see above). In supervised mode, a contrast of interest is specified (e.g. responder vs non-responder), differential expression analysis is performed using the edgeR package TMM normalization v4.4.2 (Robinson et al., 2010), followed by limma-voom (Ritchie et al., 2015), TF activity is estimated from the resulting moderated t-statistics via the consensus method, and only TFs found to be differentially active in this contrast are retained for WTCNA. In both modes, the same WTCNA, CCA cell type group analysis, and NMF latent factor computation are subsequently applied.

### Classification of response to immune checkpoint blockers inhibitors

We only considered patients treated with anti-PD-L1 or anti-PD-1. For the Gide and Mariathasan, we considered patients with ‘‘complete response’’ or ‘‘partial response’’ as responders, and patients with ‘‘progressive disease’’ or ‘‘stable disease’’ as non-responders. For the Riaz, Liu, Van Allen, Auslander (MGH) and Hugo cohorts we used the available definition of responders and non-responders.

### Machine learning for immunotherapy response prediction

To evaluate the predictive performance of CellTFusion latent factors for immunotherapy response, we used a leave-one-dataset-out (LODO) cross-validation framework implemented via pipeML v0.0.1 (Hurtado & Pancaldi, 2026), an R package that supports custom fold construction in cross-validation pipelines. This functionality allowed us to embed CellTFusion as a feature extraction step within each fold, ensuring that factor learning and projection were performed strictly within the train/test split of each iteration. In each LODO fold, one cohort was held out as the test, set while the remaining cohorts were used for training. CellTFusion was fitted on the training cohorts only, and the held-out cohort was projected into the resulting latent factor space using our projection strategy (see above), ensuring that no information from the test cohort contributed to factor learning. A classifier was trained on the training fold latent factor scores and evaluated on the projected test fold scores, with the area under the ROC curve (AUROC) as the performance metric. This procedure was repeated for each cohort, and the mean AUROC across folds was reported as the performance estimate.

After establishing predictive performance, CellTFusion was run on the full combined dataset. Feature importance was assessed using SHAP (SHapley Additive exPlanations) values (Lundberg & Lee, 2017) computed on the final trained model via the fastshap package v0.1.4 (Greenwell, 2019/2026), using Monte Carlo sampling with 1000 simulations and bias correction. SHAP values were computed across multiple resamples of the training data in parallel and summarized as the median SHAP value per sample across resamples, providing a stable estimate of each CellTFusion’s feature contribution to the target prediction. Results were visualized using the shapviz package v0.10.3 (Lundberg & Lee, 2017).

### Boruta feature selection

Relevant NMF latent factors for clinical outcome prediction were identified using the Boruta R package v9.0.0 (Kursa & Rudnicki, 2010). Boruta was run with 100 bootstrap iterations and a confirmation threshold of 90%, classifying features as confirmed, tentative, or rejected based on their importance relative to shadow features.

### Single-cell RNAseq analysis

Preprocessed single-cell RNAseq data was obtained from (Senosain et al., 2023). The Seurat package v4.3.0.1 (Hao et al., 2021) was used for downstream analysis of the data in the R environment. We performed a principal component analysis for dimensionality reduction followed by reconstruction of the neighborhood graph on the first 20 principal components, using the elbow plot, obtaining 24 clusters. Cell annotation was done with the following references: Human Primary Cell Atlas, Immune Cell Expression and Blueprint Encode using the celldex R package v1.10.1 (Aran et al., 2019). A consensus of all three annotations was taken for identification of cell-specific clusters. Pseudobulk profiles were generated using the glmGamPoi R package v1.24.0 (Ahlmann-Eltze & Huber, 2021), aggregating raw counts per sample by computing the mean expression across all cells within each sample. The resulting pseudobulk matrix was subsequently normalized to TPM, and used as input for CellTFusion.

### Cell-cell communication analysis

LIANA+ v0.1.14 (Dimitrov et al., 2024), a LIgand-receptor ANalysis framework that provides a combination of ligand-receptor methods and a collection of different databases to calculate cell-cell communication in scRNAseq data, was used to compute communication scores across cell types in the scRNA-seq data.

### Cell communication validation

To construct an undirected cell–cell communication network from LIANA+ predictions, we first aggregated results across methods using *liana_aggregate()* and filtered interactions based on statistical significance, retaining only those with an aggregate rank ≤ 0.01, as recommended in the LIANA+ documentation. Self-interactions were removed, and source–target and target–source pairs were treated as equivalent undirected edges. The frequency of each undirected cell-type pair across all retained ligand–receptor interactions was computed and used as the edge weight of the resulting LIANA+ network.

In parallel, cell groups obtained with CellTFusion were used to construct a cell-type co-occurrence network. A binary matrix encoding the presence or absence of each cell type across samples within each cell type group was derived, from which pairwise Jaccard similarity between cell types was computed and used as edge weights, producing a weighted undirected co-occurrence network in which two cell types are connected if they tend to co-occur across samples.

Both networks were restricted to their common nodes (shared cell types) prior to comparison. To assess how well CellTFusion preserves biologically relevant interactions inferred by LIANA+, two complementary metrics were computed.

***Proportion of preserved interaction frequency*:** We calculated the fraction of LIANA+’s total interaction frequency accounted for by edges shared with CellTFusion. This metric estimates the proportion of LIANA+’s total signaling weight (frequency) that is preserved by CellTFusion.

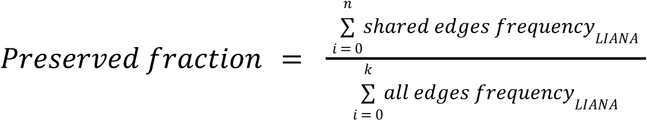

A higher value indicates that CellTFusion retains a larger portion of LIANA+’s interaction ‘volume’.

***Enrichment score for shared edges*:** To determine whether CellTFusion preferentially captures the most frequent (e.g., potentially most biologically meaningful) interactions from LIANA+, we computed an enrichment score. Specifically, we compared the mean log-transformed interaction frequency of shared edges to the mean log-transformed frequency across all LIANA+ edges:

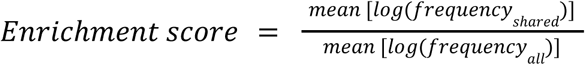

The log transformation reduces the skewness of the frequency distribution, making the comparison more robust to highly dominant edges. Values above 1 indicate that shared interactions tend to be more frequent than average in LIANA+, suggesting CellTFusion selectively recovers high-confidence communication events.

Finally, the Jaccard similarity between the two edge sets was computed as the number of shared edges divided by the total number of unique edges across the two networks, providing a symmetric measure of topological overlap independent of edge weights.

### EaSIeR immune response prediction

EaSIeR, a previously published framework for biomarker-based prediction of response to immune checkpoint blockade was used to benchmark our tool (Lapuente-Santana et al., 2021). EaSIeR integrates multiple RNA-seq–derived views of the tumor microenvironment, including immune cell fractions, pathway activities, TF activities, ligand–receptor pair weights, and cell–cell interaction scores. Starting from transcript-level TPM values, we computed established immune response signatures and EaSIeR feature views using the easier R package. Cancer-specific models trained on experimentally validated biomarkers were then applied to generate immune response predictions for each patient. Predictions were obtained independently for each molecular view and subsequently combined into an ensemble score by averaging across views. The resulting EaSIeR ensemble scores were used directly to rank patients according to predicted immune response and to evaluate predictive performance using AUROC metrics, without retraining models on the target cohorts.

### Immune heterogeneity (iHet) score

As a benchmark for immune-related tumor heterogeneity, we used the immune heterogeneity (iHet) signature previously described by (Lapuente-Santana et al., 2024). Briefly, iHet was derived using multi-omics factor analysis (MOFA) applied to bulk RNA-seq data from TCGA non-small-cell lung cancer (NSCLC) tumors, integrating tumor microenvironment cell-type composition, pathway activity, and TF activity inferred from gene expression. The first latent factor (F1), which captured the largest fraction of variance, was defined as the iHet signature. For each sample in our dataset, iHet scores were computed by projecting the normalized feature matrix onto the TCGA-derived iHet feature weights, yielding a single continuous score per sample. Higher iHet values indicate increased immune activity within the tumor microenvironment. Two additional variants of this score were evaluated alongside the base score: iHet_excl, in which weights for features positively correlated with fibroblast density are set to zero to remove the immune exclusion component, and iHet_rev, in which those weights are negated to invert the exclusion signal. The resulting iHet scores were used directly to rank samples and evaluate predictive performance without retraining or fine-tuning on the target cohort.

### Benchmarking of TME features

TME features were derived from bulk RNA-seq data and included immune activity scores, cell type deconvolution estimates, signaling pathway activities, TF activities, ligand–receptor interaction weights, and intercellular communication scores. In addition to these features, we assessed model performance using raw cell type deconvolution generated from various methods–signature combinations, as well as cell subgroups obtained using multideconv. Machine learning models were trained using each individual TME feature type and were tested on the held-out cohort to predict response to immunotherapy. Performance was evaluated using the AUROC.

### Principal component analysis (PCA)

PCA was performed on log_2_(TPM + 1) gene expression counts to assess batch effect across samples. PCA was computed using the prcomp function from the R package stats v4.4.1 and the first two principal components were retained for visualization.

### Statistical analyses

Differences in NMF factor scores between clinical groups were assessed using the Wilcoxon rank-sum test, with p-values reported for each comparison. Associations between TF module eigengenes and latent factor scores with clinical variables were quantified using Pearson correlation coefficients, with p-values derived from the asymptotic Student’s t-approximation and adjusted for multiple testing using the Bonferroni correction. For survival analysis, patients were stratified into high and low factor score groups based on the median factor score. Survival distributions were compared using the Kaplan-Meier estimator and differences between strata were assessed using the log-rank test, while hazard ratios were estimated using the Cox proportional hazards model from the R package survival v3.7.0.

## Data availability

No new experimental data was generated as part of this study. All the datasets used are publicly available.

## Code availability

The code to reproduce all the figures and analysis from this paper are available on GitHub at https://github.com/VeraPancaldiLab/CellTFusion_paper. The CellTFusion R package as well as an Rshiny application are available at https://github.com/VeraPancaldiLab/CellTFusion.

## Acknowledgements

The authors thank Leila Khajavi and Abdelmounim Essabbar for providing useful discussion on the pipeline methods and Hafida Hamdache and Francesco Massaini for testing the pipeline and providing valuable feedback. This study has been partially supported through the grant EUR CARe N°ANR-18-EURE-0003 in the framework of the Programme des Investissements d’Avenir and an Eiffel Excellence doctoral fellowship to M. H.

## Author contributions

Conceptualisation: V.P. and M.H., Methodology: M.H., Software: M.H., Validation: M.H, Formal analysis: M.H., Visualisation: M.H., R-shiny: M.H., Supervision: V.P. Writing—original draft: V.P. and M.H., Writing—review and editing: M.H and V.P.

## Additional information

**Supplementary Information.** The online version contains supplementary material available at XXX. An example tutorial and a vignette including information about the functions of the CellTFusion R package are available in the Github repository.

**Correspondence** and requests for materials should be addressed to Marcelo Hurtado or Vera Pancaldi.

