## Supplementary Material for "CellTFusion: a transcriptional regulatory network framework for the identification of functional multicellular states from bulk RNA-seq data"

### Supplementary information

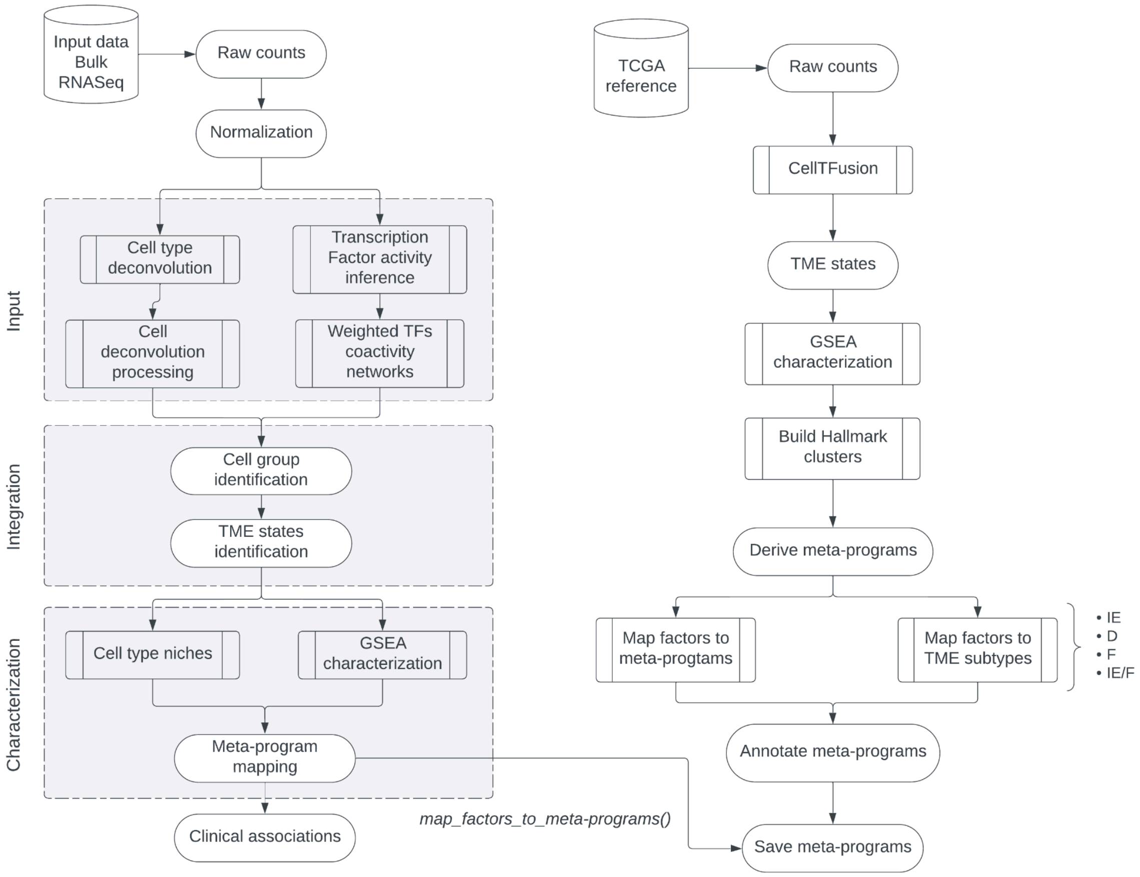


***Supplementary Figure 1.*** *Overview of the CellTFusion computational pipeline and TCGA meta-program reference construction.* ***Left:*** *CellTFusion workflow applied to a study cohort. Raw counts from bulk RNA-seq data are normalized and processed in parallel through two input streams: cell type deconvolution followed by cell deconvolution processing, and transcription factor activity inference followed by weighted TF co-activity network construction. These two streams are integrated in the integration layer, where cell groups are identified via canonical correlation analysis and compressed into TME states via non-negative matrix factorization (NMF). In the characterization layer, each TME state is annotated by its cell type niche composition and GSEA Hallmark enrichment profile, and mapped onto cancer-type-specific meta-programs.* ***Right:*** *TCGA meta-program reference construction pipeline. Raw counts from TCGA cancer-type-specific cohorts are processed through CellTFusion to derive TME states, which are characterized by GSEA and used to build Hallmark clusters. Meta-programs are derived from these clusters and annotated by TME subtype (IE: immune-enriched, D: desert, F: fibrotic, IE/F: immune-enriched fibrotic).*

*The resulting meta-programs are saved and used as a reference for mapping study cohort factors to TME subtypes.*

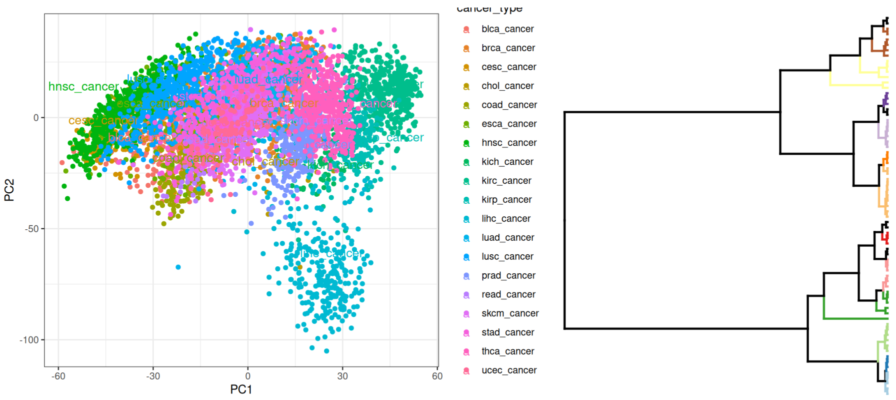


***Supplementary Figure 2. Left:*** *Principal component analysis of TCGA bulk RNA-seq data across 21 cancer types. PCA was performed on log2(TPM+1) normalized expression profiles from all TCGA samples included in the meta-program reference construction. Each dot represents one patient, colored by cancer type.* ***Right:*** *Hierarchical clustering dendrogram of cell groups identified by CellTFusion in the TCGA-SKCM cohort.*

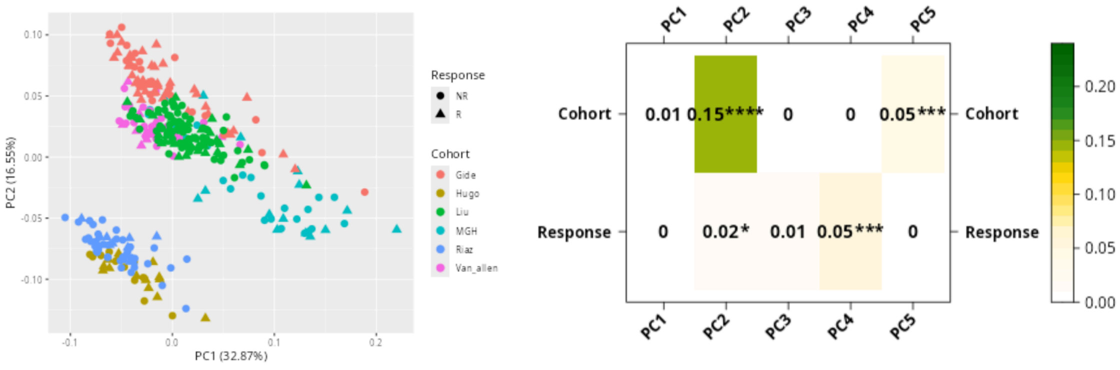


***Supplementary Figure 3. Left:*** *PCA of log-normalized gene expression across the six melanoma cohorts (Gide, Hugo, Liu, MGH, Riaz, Van_allen), colored by cohort and response status (NR: non-responder, R: responder).* ***Right:*** *Pearson correlation of principal components (PC1–PC5) with cohort and response variables. Asterisks indicate statistical significance (*p<0.05, ***p<0.001, ****p<0.0001).*

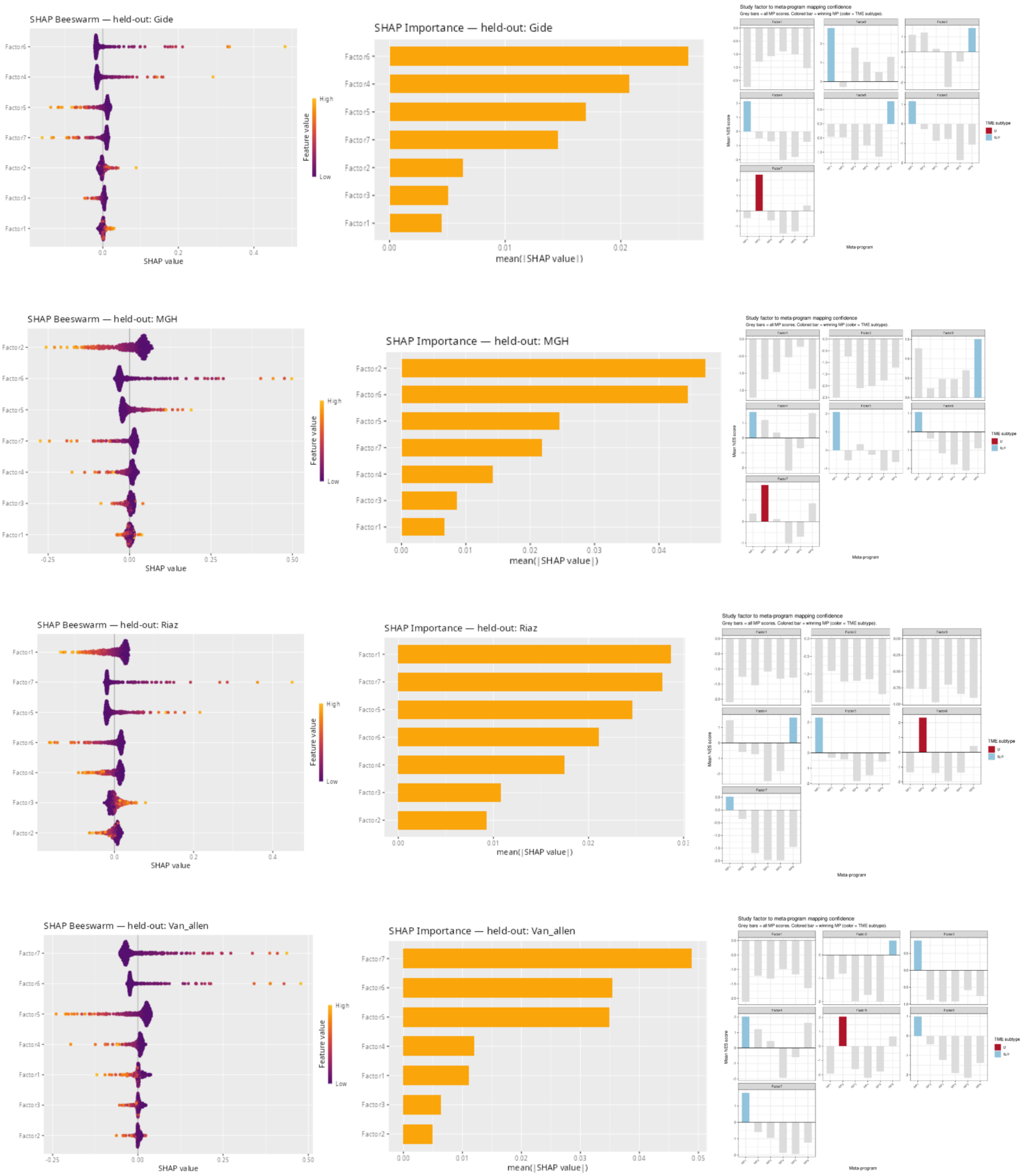


***Supplementary Figure 4. LODO fold-specific SHAP analysis and meta-program mapping for each held-out melanoma cohort.*** *For each held-out cohort (Gide, MGH, Riaz, Van_allen), three panels are shown.* ***Left:*** *SHAP beeswarm plot showing the direction and magnitude of each factor's contribution to immunotherapy response prediction; each dot represents one sample, colored by feature value (orange: high, purple: low), and the x-axis indicates the SHAP value (positive: toward response, negative: toward non-response).* ***Middle:*** *SHAP importance bar plot showing the mean absolute SHAP value for each factor,*

*reflecting the overall contribution magnitude to model predictions.* ***Right:*** *SKCM meta-program mapping confidence for the corresponding LODO training partition. Grey bars show NES scores against all meta-programs (MP1–MP6); the colored bar indicates the winning meta-program, colored by TME subtype (blue: IE/F, red: D). Since CellTFusion is independently re-run for each training partition, factor indices are not comparable across cohorts and should be interpreted with their meta-program assignment shown in the right panel.*

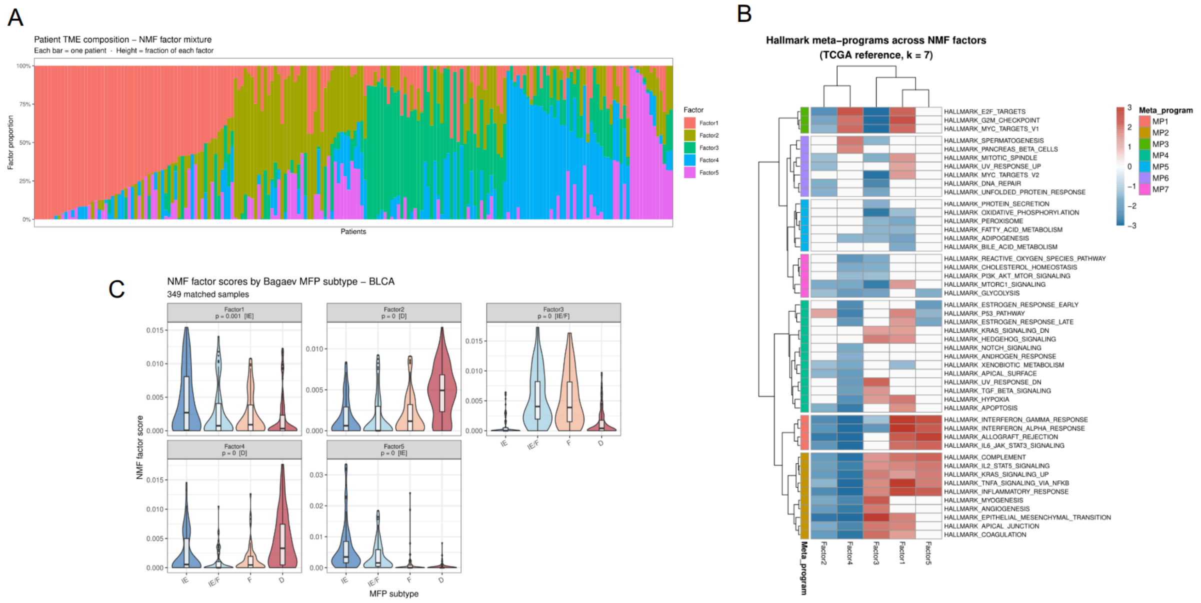


***Supplementary Figure 5. TCGA-derived transcriptional meta-program for BCLA. (A)*** *Patient TME composition barplot for BCLA (n=349). Each bar represents one patient, colored by NMF factor proportion (Factor1, Factor2, Factor3, Factor4, Factor5). Patients are ordered by dominant factors.* ***(B)*** *Hallmark meta-program heatmap for BCLA (k=7). Rows are MSigDB Hallmark gene sets, columns are NMF factors, and color represents the normalized enrichment score (NES). Hallmarks are grouped into seven meta-programs (MP1–MP7).* ***(C)*** *NMF factor scores stratified by Bagaev et al., 2021 into Molecular Functional Phenotype (MFP) subtype for BCLA. Violin plots show factor score distributions across IE, IE/F, F, and D subtype groups. Kruskal-Wallis p-values are indicated.*

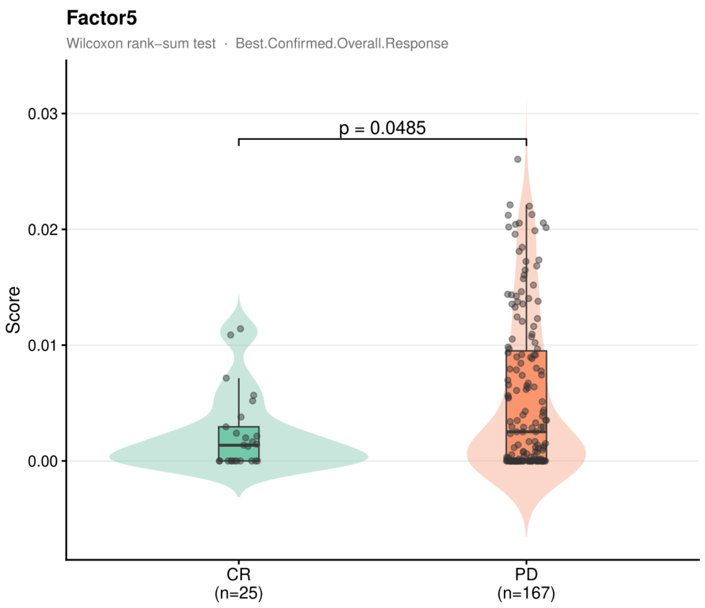


***Supplementary Figure 6. Pairwise association of CellTFusion factor scores with immunotherapy response in the bladder cancer cohort.*** *Violin plots show the distribution of Factor 5 scores stratified by best confirmed overall response (CR: complete response, n=25; PD: progressive disease, n=167). Wilcoxon rank-sum test p-value is indicated. Factor 5 showed a nominally significant difference (p = 0.049) with higher scores in PD patients.*

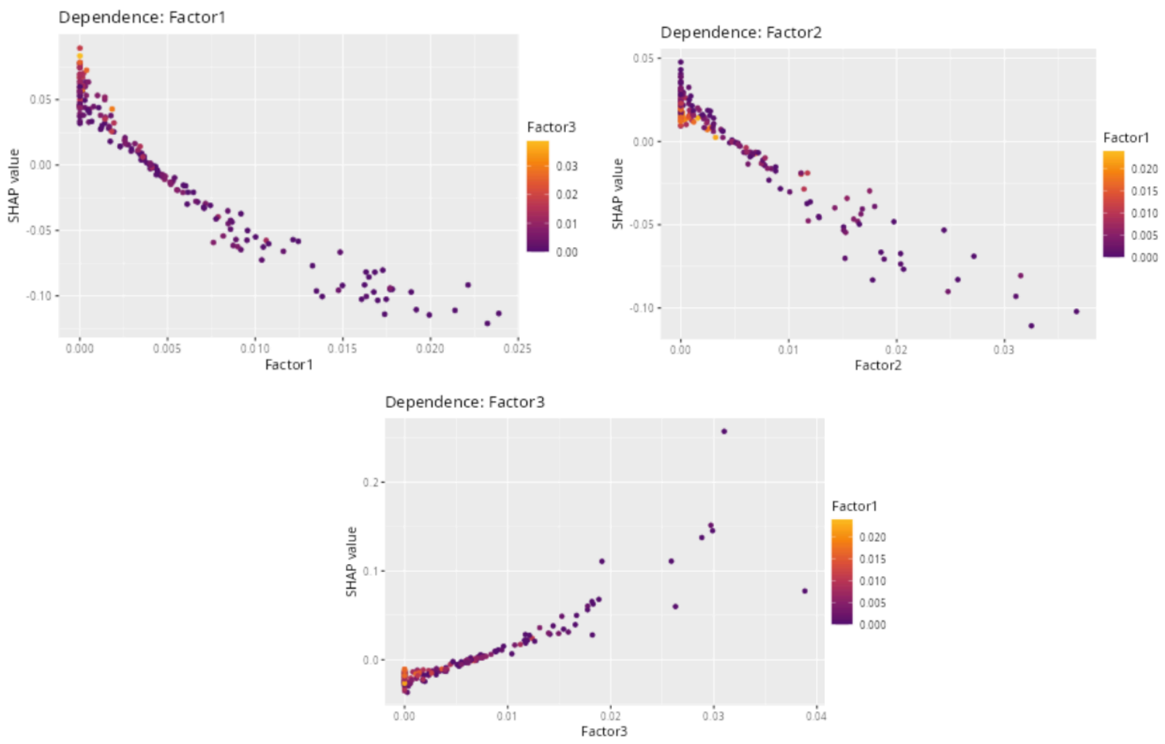

***Supplementary Figure 7. SHAP dependence plots for the top contributing factors in the bladder cancer cohort.*** *Each panel shows the relationship between a factor's score (x-axis) and its SHAP value (y-axis), with dots colored by the score of the most interacting feature. Factor 1 and Factor 2 show a clear negative dependence, where increasing factor scores are associated with decreasing SHAP values, indicating that high scores of both factors push predictions toward progressive disease. Factor 3 shows a positive dependence, where increasing scores are associated with increasing SHAP values, confirming that high Factor 3 scores push predictions toward complete response.*

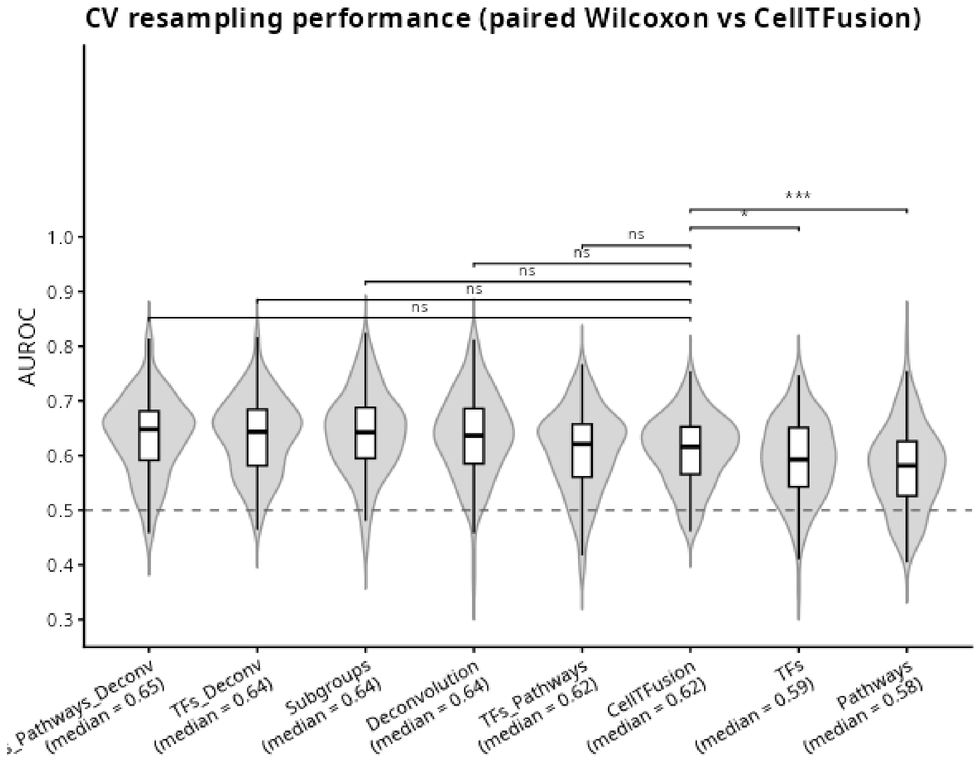


***Supplementary Figure 8. Within-cohort cross-validation resampling performance comparison across TME feature frameworks.*** *Violin plots show the distribution of AUROC values across resampling iterations for each feature framework in the combined melanoma cohort. Median AUROC values are indicated for each framework: TFs + Pathways + Deconvolution (0.65), TFs + Deconvolution (0.64), Subgroups (0.64), Deconvolution (0.64), TFs + Pathways (0.62), CellTFusion (0.62), TFs (0.59), and Pathways (0.58). Brackets indicate pairwise comparisons against CellTFusion assessed by paired Wilcoxon test (ns: not significant, *: $p < 0.05$, ***: $p < 0.001$). The dashed line indicates random classifier performance (AUROC = 0.5).*
